# Hmgb2 improves astrocyte to neuron conversion by increasing the chromatin accessibility of genes associated with neuronal maturation in a proneuronal factor-dependent manner

**DOI:** 10.1101/2023.08.31.555708

**Authors:** Priya Maddhesiya, Tjasa Lepko, Andrea Steiner-Mezzardi, Veronika Schwarz, Juliane Merl-Pham, Finja Berger, Stefanie M. Hauck, Lorenza Ronfani, Marco Bianchi, Giacomo Masserdotti, Magdalena Götz, Jovica Ninkovic

## Abstract

**Background:** Direct conversion of reactive glial cells to neurons is a promising avenue for neuronal replacement therapies after brain injury or neurodegeneration. The overexpression of neurogenic fate determinants in glial cells results in conversion to neurons. For repair purposes, the conversion should ideally be induced in the pathology-induced neuroinflammatory environment. However, very little is known regarding the influence of the injury-induced neuroinflammatory environment and released growth factors on the direct conversion process.

**Results:** We established a new *in vitro* culture system of postnatal astrocytes without epidermal growth factor that reflects the direct conversion rate in the injured, neuroinflammatory environment *in vivo*. We demonstrated that the growth factor combination corresponding to the injured environment defines the ability of glia to be directly converted to neurons. Using this culture system, we showed that chromatin structural protein high mobility group box 2 (HMGB2) regulates the direct conversion rate downstream of the growth factor combination. We further demonstrated that Hmgb2 cooperates with neurogenic fate determinants, such as Neurog2, in opening chromatin at the loci of genes regulating neuronal maturation and synapse formation. Consequently, early chromatin rearrangements occur during direct fate conversion and are necessary for full fate conversion.

**Conclusions:** Our data demonstrate novel growth factor-controlled regulation of gene expression during direct fate conversion. This regulation is crucial for proper maturation of induced neurons and could be targeted to improve the repair process.

## Background

Innovative approaches to stimulate tissue regeneration and functional restoration of the central nervous system are required, because the adult mammalian brain has limited ability to replace lost neurons [1–4]. Direct conversion of glial cells to neurons (induced neurons, iN) is a promising avenue for successful repair [2,5,6]. The overexpression of several neurogenic factors, alone or in combination, induces the conversion of several cell types, including astrocytes, pericytes, oligodendrocyte progenitors and fibroblasts, into post-mitotic neurons with different well-defined neurotransmitter identities [7–24]. These strong inducers of the neurogenic fate are transcription factors (TFs) that specify neuronal fate during development [7]. Many of these TFs have recently been shown to have pioneering factor activity and to bind closed chromatin configurations [5,25,26]. Indeed, recent insights regarding the fundamentals of neuronal fate specification have revealed that changes in chromatin structure might be a key factor in the stable acquisition of neuronal fate [27,28], in line with the pioneering activity of fate determinants inducing fate conversion. Despite their remarkable strength, defined single pioneering TFs (e.g., Neurog2) cannot successfully reprogram some starting cell types or cell states induced by culturing conditions [14]. The inability of Neurog2 to activate gene expression has been associated with epigenetic silencing of target loci [14,29]. Interestingly, forskolin (an agonist of adenylyl cyclase) and dorsomorphin (an inhibitor of BMP signaling) enhance the chromatin accessibility mediated by Neurog2, thus suggesting that additional pathways contribute to Neurog2’s trailblazing properties [30,31]. In fact, treating Neurog2-expressing cells with these small molecules results in chromatin opening at a substantial number of sites, including CRE half-sites or HMG box motifs [30]. Thus, small molecules or a combination of other TFs may be necessary to induce successful or efficient reprogramming, depending on the starting populations, although Neurog2 is a pioneer factor that can overcome the lineage barrier. In addition to several factors associated with chromatin, microRNAs and small molecules have been found to improve the conversion efficiency and maturation status of reprogrammed neurons despite being unable to induce conversion on their own [12,15,32,33]. These findings support a model in which multi-level lineage barriers maintain cell identity and must be overcome for cells to acquire neuronal fate adequate for repair purposes. Comprehensive understanding of these barriers is at the core of successful iN generation and the functional restoration of the damaged CNS.

Importantly, most of these barriers have been identified through the use of defined and stable *in vitro* systems. However, for repair purposes, iNs must be generated in the injured environment. The intricacy of the injured milieu is an obstacle to understanding the molecular mechanisms of direct neuronal conversion *in vivo*. Injury triggers the release of several signaling factors with precise temporal resolution that can either resolve or strengthen the lineage barriers [34]. For example, epidermal growth factor (EGF) levels spike within 24 hours after brain injury and remain elevated for 3 days before returning to baseline. In contrast, basic fibroblast growth factor (bFGF) levels begin to rise 4 hours after damage and remain elevated for 14 days [34]. Infusion of bFGF into the brain after traumatic brain injury, for example, greatly enhances cognitive performance in animals by increasing neurogenesis [35]. Additionally, EGF infusion enhances neurogenesis via enlargement of the neurogenic precursor pool in the neurogenic niche after ischemia injury [36]. Moreover, forced Neurog2 expression in glial cells, along with the bFGF2 and EGF growth factors, enhances neuronal reprogramming *in vivo* [37]. Importantly, EGF receptor (EGFR) signaling has been proposed to regulate both global chromatin state and the accessibility of specific loci [38]. Furthermore, interaction of EGFR signaling and chromatin remodelers from the SWI/SNF family is critical for the expansion of beta cells after pancreas injury [39]. Similarly, FGF signaling orchestrates chromatin organization during neuronal differentiation [40]. Together, environmental signals are likely to be integrated into the lineage barriers defining the propensity of starting glial cells to be converted to postmitotic neurons.

To investigate the embedding of growth factors in lineage barriers relevant to *in vivo* direct neuronal reprogramming after brain injury, we developed an *in vitro* model with altered growth factor composition. We showed that, in this model, neurogenic fate determinants induced astrocyte to neuron conversion with a diminished efficiency comparable to the conversion rate observed *in vivo.* This system allowed us to identify Hmgb2 as a novel regulator in the context of direct astrocyte to neuron conversion. We showed that high levels of Hmgb2 alleviate the lineage barrier and promote efficient establishment of neuronal fate. Our data suggest that Hmgb2-dependent chromatin opening of regulatory elements controls the expression of neuronal maturation genes and enables the establishment of the full neurogenic program, thereby resulting in efficient astrocyte to neuron conversion.

## Results

### Growth factors shape the lineage barriers to glia to neuron conversion

To investigate the contributions of injury-induced growth factors to lineage barriers to maintaining glial fate in the injured mammalian brain, we established a new *in vitro* model with the growth factor composition adjusted to better reflect the local environment after injury. After brain injury, levels of EGF peak within the first 24 h and return to baseline levels 3 days post injury (dpi). In contrast, FGF levels increase by 4 h after injury and persist until 14 dpi [41]. To mimic the dynamics in the *in vivo* environment, we cultured astrocytes, obtained from postnatal murine cerebral cortex (P5–P7) for 10 days in the presence of only bFGF, then compared the direct conversion rates to neurons in this culture with the conversion efficiency in the widely used culture conditions containing both EGF and bFGF [42,43]. To convert astrocytes into neurons, we transduced cells with an MLV-based retrovirus for expression of the neurogenic TFs reported to reprogram astrocytes (Neurog2, Pou3f2 or Sox11; Fig. 1a) *in vitro* and a fluorescent reporter protein. The expression of the fluorescent reporter protein was used to identify the transduced cells. The identity of the transduced cells was probed 7 days after viral transduction (days *in vitro* (div); Fig. 1a). Only cells expressing doublecortin (DCX) and having at least one process longer than three cell somata diameters were identified as neuronal cells, according to Gascon et al. [44] (Fig. 1b, c). The transduction of astrocytes with control viruses for expression of either GFP or dsRed did not induce glia to neuron conversion in any culturing conditions (Suppl. Fig. 1a-d). In contrast, the transduction of astrocytes isolated from EGF+bFGF culture with several neurogenic fate determinants did induce their conversion, and neurons at different maturation stages (on the basis of the complexity of their processes) were observed after 7 div (Fig. 1b, d). Interestingly, neither Neurog2 nor Pou3f2 induced the direct conversion of astrocytes grown in the presence of only bFGF, whereas the culturing conditions did not significantly alter the conversion by overexpression of Sox 11 (Fig. 1d). Because the culture condition with bFGF contained only half the usual growth factors, we assessed the conversion rate of cultures containing only EGF. Importantly, Neurog2 induced the conversion of astrocytes grown with only EGF at the same rate as astrocytes grown in EGF+bFGF culture medium (Suppl. Fig. 1d-f), in line with the specific role of bFGF in decreasing the conversion rate.

**Figure 1.**
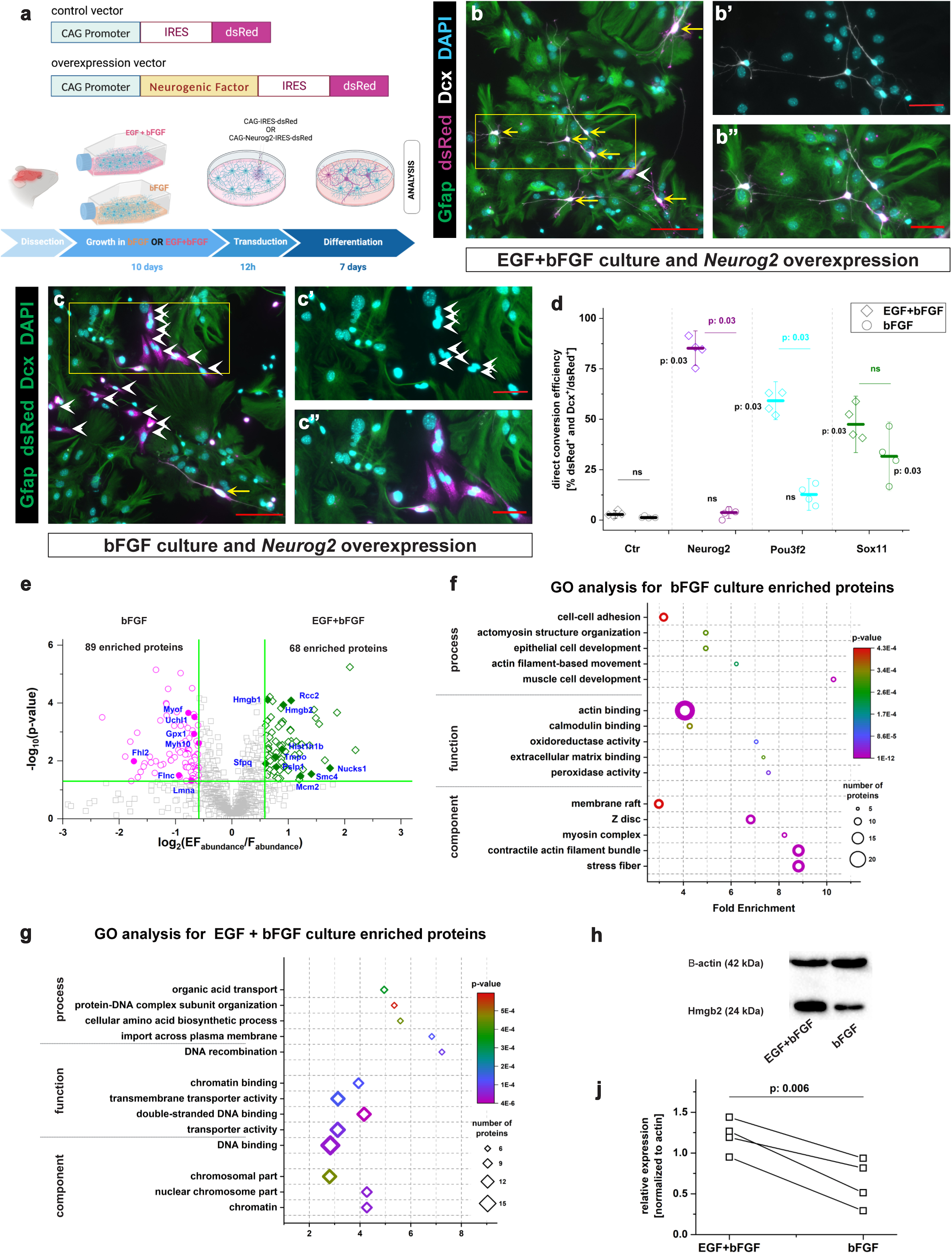
Astrocyte growth conditions define the rate of direct astrocyte to neuron conversion. (**a**) Schemes depicting viral vector design and the experimental paradigm used for astrocyte to neuron conversion. (**b-c’’**) Micrographs illustrating the identity of Neurog-Neurog2 transduced cells 7 days after transduction in the EGF+bFGF (b) and bFGF (c) culture conditions. b’, b’’, c’ and c’ are magnifications of boxed areas in b and c, respectively. Yellow arrows indicate successfully converted cells, whereas white arrowheads indicate cells failing to convert. Scale bars: 100 µm in b and c; 50 µm in b’, b’’, c’ and c’’. (**d**) Dot plot depicting the proportion of transduced cells converting to neurons in EGF+bFGF and bFGF cultures 7 days after transduction with different neurogenic fate determinants. Data are shown as median±IQR; each single dot represents an independent biological replicate. Significance was tested with two-tailed Mann-Whitney test. p-values: black font corresponds to the comparison to the control and colored to the comparison between EGF+bFGF and bFGF. (**e**) Volcano plot depicting proteins enriched in astrocytes cultured in bFGF (magenta circles) and EGF+bFGF (green diamonds) culture conditions (fold change >1,5; p value <0,05). (**f**, **g**) Plots depicting the top five enriched GO terms in protein sets enriched in bFGF (f) and EGF+bFGF (g) cultures. (**h**) Western blot depicting levels of Hmgb2 protein in EGF+bFGF and bFGF astrocyte cultures. (**j**) Dot plot showing the relative levels of Hmgb2 (normalized to actin) in EGF+bFGF and bFGF cultures. Data are shown as median±IQR; single dots represent independent biological replicates. Paired-t-test was used for the significance test. Abbreviation: GO, Gene Ontology.

This difference in direct conversion could be explained by the selection of particular cell types during astrocyte expansion with growth factors. Therefore, we assessed the identity of the transduced cells 24 h after transduction by using immunocytochemistry (Suppl. Fig. 2a). Most cells expressed the astrocyte marker S100β in both culture conditions, without any significant differences (Suppl. Fig. 2b, c, f). Similarly, we did not observe any differences in the proportion of GFAP+ cells (Suppl. Fig. 2d-f). In line with reports that astrocytes *in vitro* express the TF Olig2 [45], most cells in both culture conditions expressed Olig2 (Suppl. Fig. 2g-i). Moreover, we observed only a small proportion of DCX+ neuronal progenitors or αSMA+ pericytes in both cultures (Suppl. Fig. 2d-i), thus indicating comparable cellular compositions between cultures, according to the analyzed marker expression. Interestingly, we observed lower proliferation rates of astrocytes grown in bFGF than EGF+bFGF conditions, on the basis of the expression of Ki67 or pH3 (Suppl. Fig. 2j-n). This finding suggested that bFGF-grown astrocytes might further differentiate, epigenetically silence neuronal loci and become less prone to direct conversion, as previously shown for long-term astrocyte cultures [46]. To examine this possibility, we cultured astrocytes for 7 days in bFGF culture conditions, added EGF and grew astrocytes for an additional 7 days with EGF+bFGF (Suppl. Fig 3a). The conversion rate of these astrocytes was compared with that of astrocytes cultured in either EGF+bFGF or bFGF for 14 days (Suppl. Fig. 3b, f). As expected, longer culturing of cells in either bFGF or EGF+bFGF decreased the direct reprogramming rate (Suppl. Fig. f), as previously described [46]. However, the post-culturing of initially bFGF-grown astrocytes in EGF+bFGF for 7 days improved their reprogrammability, and we observed no differences in the proportions of generated neurons compared with astrocytes continuously cultured in EGF+bFGF (Suppl. Fig. 3b, c, f). Moreover, the conversion rate of EGF+bFGF-grown astrocytes decreased after culturing in bFGF for 7 days, and no differences were observed between this culture and continuously bFGF cultured astrocytes (Suppl. Fig. 3d-f). Together, the cell identity marker analysis and the alterations in the culture composition experiments suggested that growth factor conditions define the astrocytic lineage barriers and consequently the rate of direct conversion to neurons, on the basis of neurogenic factor overexpression.

### High mobility group box 2 (Hmgb2) levels are decreased in bFGF astrocyte culture

To identify factors responsible for maintaining the astrocytic lineage barrier, we performed label-free LC-MS/MS-based proteome analysis of astrocytes cultured with either bFGF or EGF+bFGF for 10 days. In total, we detected approximately 1700 proteins, of which 157 showed differences in levels between culture conditions (1.5-fold change, p<0.05): 68 significantly enriched in the EGF+bFGF culture and 89 significantly enriched in the bFGF culture (Fig. 1e, Suppl. Table 1). Gene Ontology (GO) analysis revealed an enrichment of cytoskeleton-associated processes in the protein set enriched in the bFGF-grown culture (Fig. 1f; Suppl. Table 1), whereas transport across the mitochondrial membrane, metabolic processes and chromatin-associated processes were enriched in the EFG+bFGF induced proteome (2-fold enrichment, p<0.05; Fig. 1g). These data are in line with recent evidence indicating that changes in the mitochondrial proteome during astroglia to neuron conversion determine the extent of the direct conversion [47]. Moreover, because chromatin state has been reported to regulate lineage barriers in reprogramming [44,48–52], we searched for chromatin-associated factors differentially enriched between culture systems. The chromatin architectural protein Hmgb2 was 1,88-fold enriched in EGF+bFGF compared with bFGF cultures (Fig. 1e). This enrichment was confirmed by western blotting (Fig. 1h, j). Interestingly, we also observed that the HMGB2 protein family member HMGB1 was enriched in the EGF+bFGF culture condition, although at a lower level (Fig. 1e). In the adult mouse brain, Hmgb2 is specifically expressed in cells committed to the neurogenic lineage (transit amplifying progenitors, neuroblasts) in both neurogenic niches [53] in addition, traumatic brain injury induces Hmgb2 expression in a subset of reactive astrocytes (Suppl. Fig. 4). These findings suggest that HMGB2 might be an important factor improving direct conversion in the EGF+bFGF culture.

### Hmgb2 levels define the rate of direct astrocyte to neuron conversion

To test whether Hmgb2 might have functional relevance in fate conversion, we transduced astrocytes, grown for 10 days in medium containing either EGF+bFGF or bFGF, with Hmgb2-encoding retrovirus (Fig. 2a), and assessed the identity of the transduced cells 7 days later, on the basis of DCX expression and cell morphology (see above; Fig. 1b-d). Overexpression of Hmgb2 did not alter cell identity in either culture condition (Fig. 2b-e). Most cells retained their astrocyte identity and expressed GFAP (Fig. 2e). However, when we co-transduced the bFGF-grown astrocytes with retroviruses for expression of Neurog2-dsRED and Hmgb2-GFP, we observed a 2.5-fold greater conversion rate in the co-transduced cells than cells transduced with Neurog2 only (Fig. 2c, d). Interestingly, the co-overexpression of Neurog2+Hmgb2 did not further improve the conversion of EGF+bFGF-grown astrocytes, because the conversion rate of Neurog2+Hmgb2 co-transduced astrocytes was comparable to that of Neurog2-transduced astrocytes in this culture condition (Fig. 2b, d).

**Figure 2.**
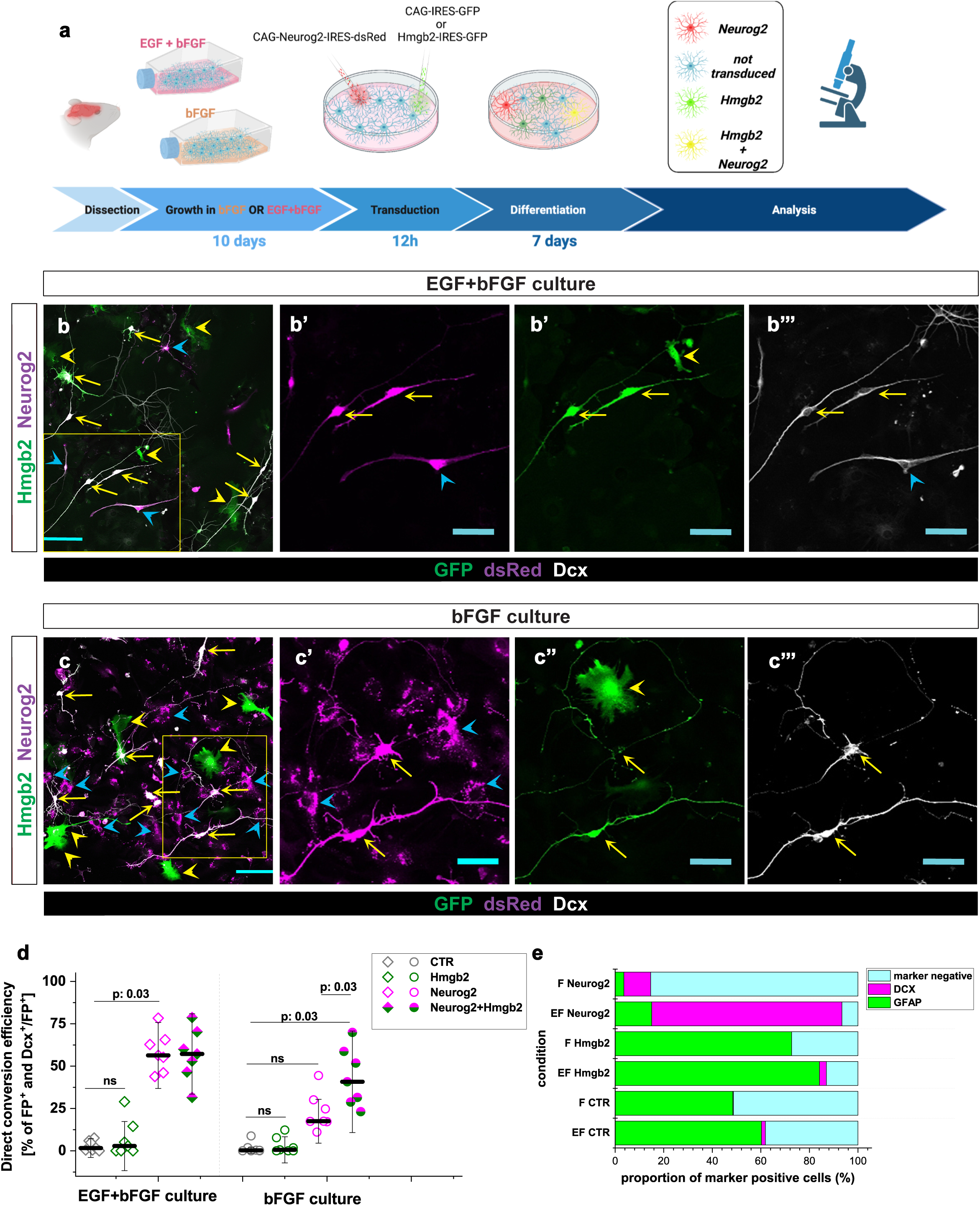
Hmgb2 is sufficient for successful Neurog2-mediated direct astrocyte to neuron conversion. (**a**) Scheme depicting the experimental paradigm used for astrocyte to neuron conversion. (**b-c’’’**) Micrographs showing the identity of Neurog2- and Hmgb2-expressing virally transduced cells 7 days after transduction in EGF+bFGF (a) and bFGF cultures (b). b’, b’’, b’’’, c’, c’’ and c’’’ are magnifications of the boxed areas in a and b, respectively. Yellow arrows indicate co-transduced cells expressing Neurog2 and Hmgb2, yellow arrowheads indicate cells transduced only with Hmgb2-encoding virus, and blue arrowheads indicate cells transduced with only Neurog2-encoding virus. Scale bars: 100 µm in b and c; 50 µm in b’. b’’, b’’’, c’, c’’ and cb’’’. (**d**) Dot plot depicting the proportion of transduced cells converting to neurons in EGF+bFGF and bFGF cultures 7 days after transduction. Data are shown as median±IQR; single dots represent independent biological replicates. Significance was tested with two-tailed Mann-Whitney test. (**e**) Histogram depicting the identities of cells transduced with the indicated factors 7 days after transduction. Abbreviation: FP, fluorescent protein.

Improvement in the Neurog2-mediated conversion rate of bFGF-grown astrocytes prompted us to investigate whether this improvement might be factor-specific. Therefore, we assessed the effect of Hmgb2 overexpression on Pou3f2-mediated fate conversion, given that the neurogenic capability of Pou3f2 was also diminished in bFGF-grown astrocytes (Fig. 1d). Similarly to the Neurog2-mediated conversion, the simultaneous overexpression of Hmgb2 and Pou3f2 in EGF+bFGF-grown astrocytes did not result in higher conversion rates, whereas the factor combination significantly increased the conversion rate in bFGF-grown astrocytes (Suppl. Fig. 1g). Together, these data suggested that Hmgb2 does not induce direct conversion on its own but increases the ability of neurogenic factors to overcome the lineage barriers.

To test whether Hmgb2 might be necessary for direct astrocyte to neuron conversion, we isolated astroglia from Hmgb2-deficient mice (Hmgb2^MUT/MUT^) and their siblings (Hmgb2^WT/MUT^ and Hmgb2^WT/WT^), cultured them in the direct conversion permissive conditions (EGF+bFGF) and induced conversion by Neurog2 overexpression (Fig. 3a). Neurog2 overexpression induced direct conversion of Hmgb2^WT/WT^ and Hmgb2^WT/MUT^ astrocytes (Fig. 3b-d), in agreement with our previous findings demonstrating high responsiveness of EGF+bFGF-grown astrocytes (Fig. 1d). However, the conversion rate of Hmgb2-deficient (Hmgb2^MUT/MUT^) astroglia significantly decreased compared to WT siblings (Fig. 3c, d). These findings supported our hypothesis that Hmgb2 levels define the astrocytic lineage barrier.

**Figure 3.**
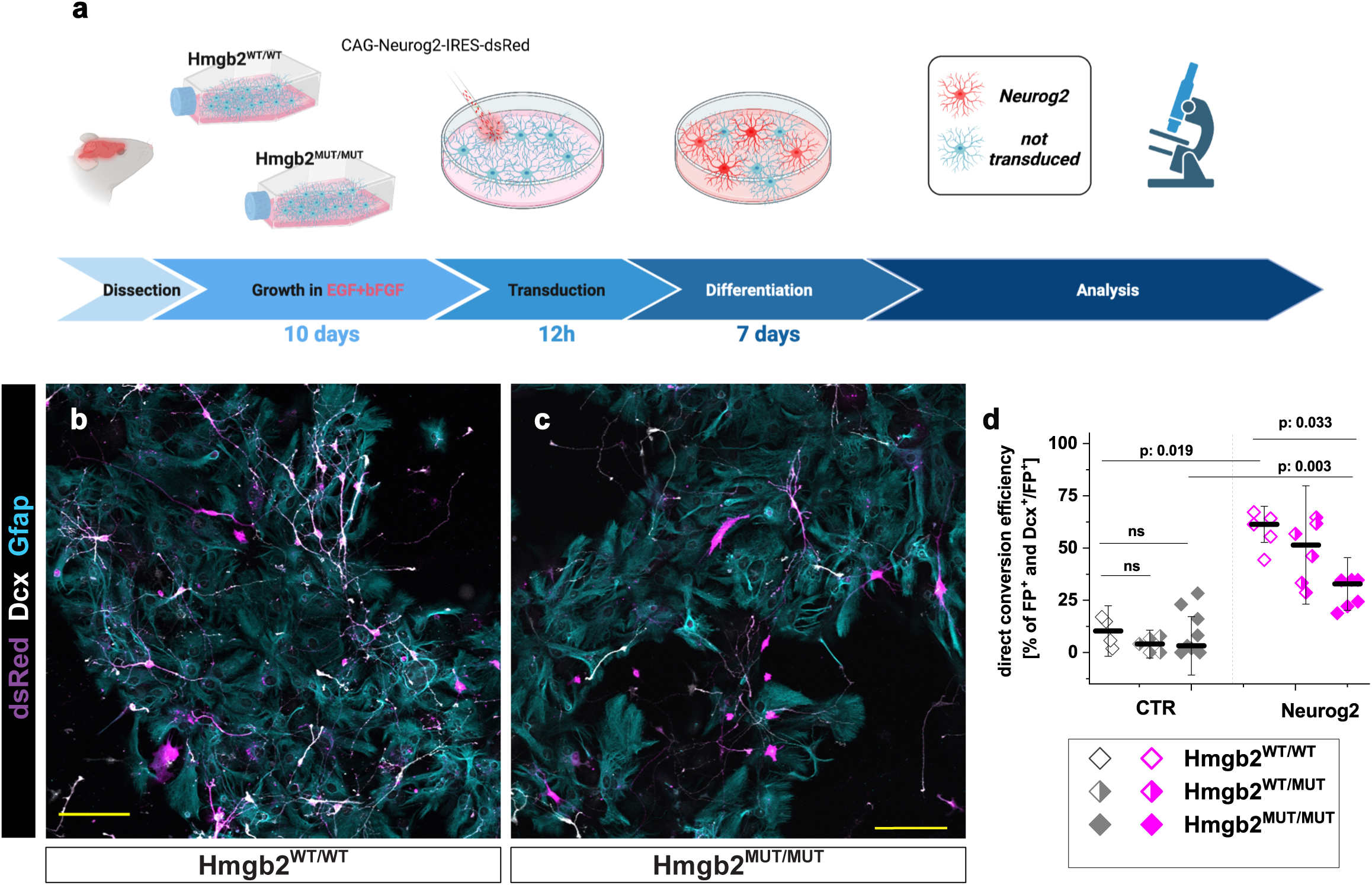
Hmgb2 is necessary for successful Neurog2-mediated direct astrocyte to neuron conversion. (**a**) Scheme depicting the experimental paradigm used for astrocyte to neuron conversion. (**b, c**) Micrographs showing the identities of Neurog2-expressing virally transduced cells 7 days after transduction in EGF+bFGF culture of astrocytes derived from Hmgb2-deficient animals (c) and their siblings (b). Scale bars: 100 µm. (**d**) Dot plot depicting the proportion of Hmgb2-deficient or control cells converting to neurons 7 days after transduction with Neurog2. Data are shown as median±IQR; single dots represent independent biological replicates. Significance was tested with two-tailed Mann-Whitney test. Abbreviation: FP, fluorescent protein.

### Prospero homeobox protein 1 (Prox1) overexpression improves direct glia to neuron conversion in FGF only culture

To understand the Hmgb2-dependent lineage barrier in direct glia to neuron conversion, we compared the transcriptional changes induced by Neurog2 overexpression in the bFGF and EGF+bFGF cultured cells 48 h after transduction. Cells transduced with different viruses were purified by FACS, and genes regulated by Neurog2 overexpression were compared (Suppl. Fig. 5). We identified differences in the expression of 443 genes (321 up-regulated and 122 down-regulated genes, fold change > 2, padj < 0.05) induced by Neurog2, as compared with that in control CAG-GFP virally transduced cells in the EGF+bFGF culture condition (Suppl. Fig. 6 a, Suppl. Table 2). In the bFGF culture, Neurog2, as compared with the respective CAG-GFP transduced control, induced 171 genes (137 up-regulated and 34 down-regulated genes, fold change > 2, padj < 0.05) (Suppl. Fig. 6 b, Suppl. Table 2). GO analysis (biological processes, fold enrichment > 2 and p < 0.05) of genes (321) upregulated in EGF+bFGF culture revealed enrichment in the terms nervous system development, neuronal differentiation, and migration (Fig. 4a), in line with the ability of Neurog2 to successfully convert astroglia to neurons. Unexpectedly, the significantly enriched biological processes in the set of the 137 up-regulated genes in the bFGF culture were also associated with regulation of neurogenesis, nervous system development and synaptic signaling (Fig. 4b), thereby indicating that Neurog2 overexpression at least partially induced the neuronal fate in astrocytes grown in the bFGF condition. Indeed, we observed that 96 genes were induced by Neurog2 in both bFGF and EGF+bFGF cultures (Fig. 4c), and were enriched in GO biological processes associated with regulation of neurogenesis, nervous system development, neuronal differentiation and migration (Suppl. Fig. 6c). In addition, in the bFGF culture, the 41 genes uniquely induced by Neurog2 (Fig. 4c) were associated with GO biological processes of cardiac muscle tissue development, leukocyte differentiation, response to lithium-ion and neurotransmitter receptor to the plasma membrane (Suppl. Fig. 6d). These findings suggested that, in contrast to the EGF+bFGF culture, in the bFGF culture, Neurog2 induced other fates along with neuronal processes possibly interfering with the establishment of the neuronal identity [54]. Furthermore, we identified 225 uniquely Neurog2-induced genes in the EGF+bFGF culture (Fig. 4c) associated with the GO biological processes regulation of membrane potential and ephrin receptor pathway (Suppl. Fig. 6d), which regulate neuronal maturation and axonogenesis [55,56]. Moreover, previously reported Neurog2-induced genes necessary for successful conversion, such as *Neurod4*, *Insm1*, *Hes6*, *Slit1*, *Sox11* and *Gang4* [46] were up-regulated in both cultures (Fig. 4d). Nevertheless, genes such as *Dscaml1*, *Prox1*, *Lrp8* and *Shf* were induced in only the EGF+bFGF culture. Importantly, the co-expression of Neurog2 and Hmgb2 in bFGF-grown astrocytes induced the expression of these genes to levels similar to those detected in the Neurog2-transduced EGF+bFGF culture (Fig. 4d). Therefore, the bFGF culture established the lineage barrier by interfering with the induction of a small, specific set of genes relevant for the conversion. To test this hypothesis, we selected one candidate, Prox1, and evaluated whether it might help overcome the bFGF only medium restrictive conditions. We overexpressed Prox1 in the bFGF-cultured cells and observed only a small increase in the conversion rate (Fig. 4e). However, after the co-expression of Neurog2 and Prox1 in bFGF-cultured astrocytes, we observed a significant increase in the proportion of generated neurons similar to the conversion rate induced by Neurog2 in the EGF+bFGF culture and the bFGF-cultured astrocytes co-transduced with Neurog2 and Hmgb2 (Fig. 4e). Moreover, microRNA-mediated knockdown of Prox1 decreased the Neurog2-mediated conversion of EGF+bFGF cultured astrocytes, in line with previous reports [46]. This conversion rate was also comparable to the rate of Neurog2-mediated conversion of bFGF-cultured astrocytes (Fig. 4e).

**Figure 4.**
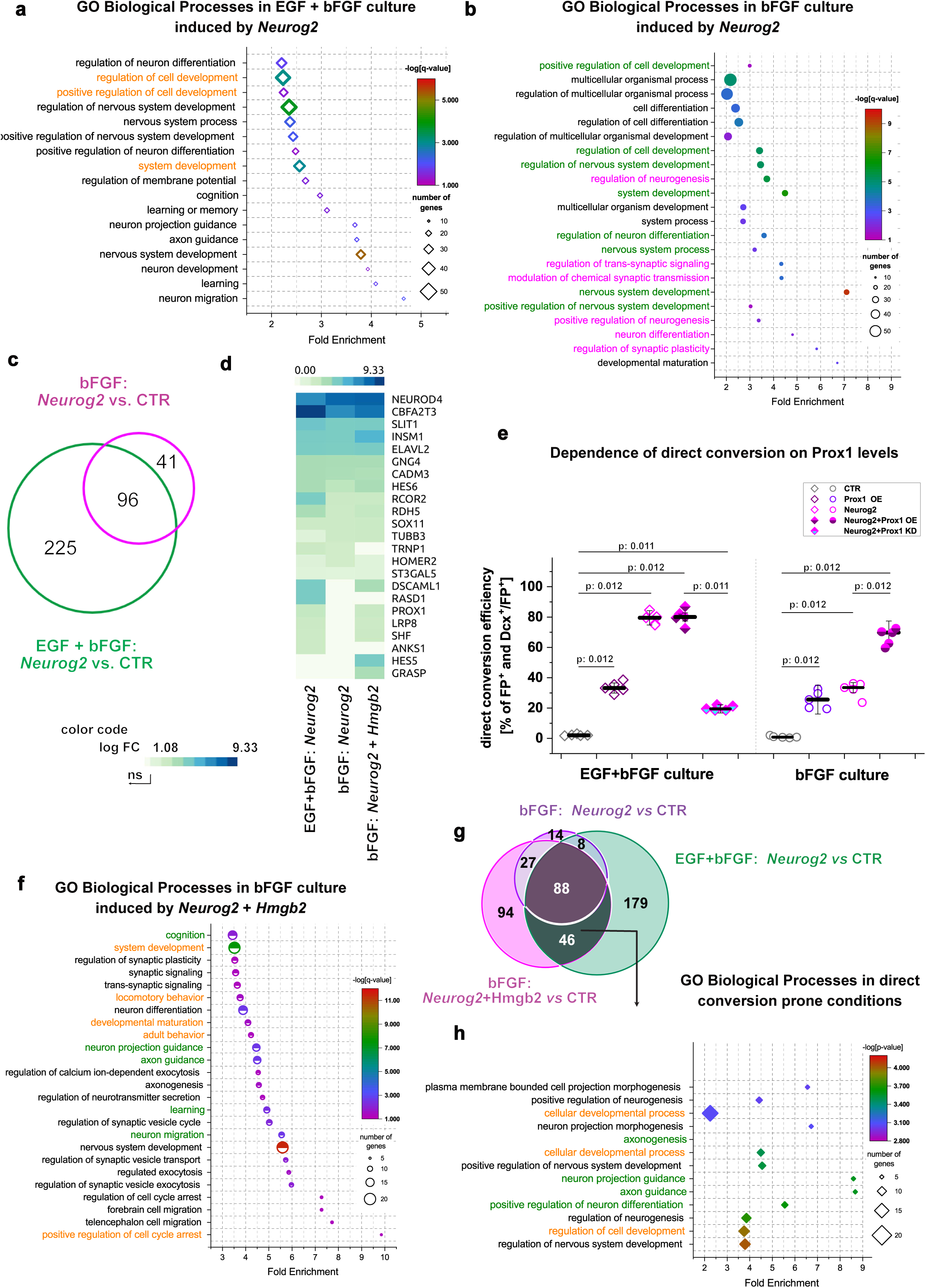
Neurog2 induces incomplete neuronal fate in bFGF culture. (**a, b**) Plots depicting enriched GO biological process terms in gene sets induced by Neurog2 in EGF+bFGF culture (a) and bFGF culture (b) 48 hours after viral transduction. Orange text represents the GO terms not associated with neuronal fate. Green and magenta text represent GO terms specifically enriched in EGF+bFGF culture and bFGF culture, respectively. (**c**) Venn diagram illustrating the overlap of Neurog2-induced transcripts in EGF+bFGF and bFGF culture 48 h after viral transduction. (**d**) Heat map showing Neurog2- or Neurog2+HMGB2-mediated induction of core neurogenic factors (according to Masserdotti et al., 2013) in EGF+bFGF and bFGF cultures. (**e**) Dot plot depicting the proportion of transduced cells converting to neurons in EGF+bFGF and bFGF cultures 7 days after transduction in Prox1 deficient or Prox-1 overexpressing cells. Data are shown as median±IQR; single dots represent independent biological replicates. Significance was tested with two-tailed Mann-Whitney test. (**f**) Plot showing GO terms enriched in the gene set upregulated in bFGF culture by Neurog2 and Hmgb2 expression 48 h after viral transduction. GO terms in green text are also induced by Neurog2 alone in EGF+bFGF culture (panel a). (**g**) Venn diagram illustrating the overlap of Neurog2-induced transcripts in EGF+bFGF and bFGF culture with Neurog2 and Hmgb2-induced transcripts after overexpression in bFGF culture 48 h after viral transduction. (**h**) Plot depicting enriched GO biological process terms in gene sets induced in the reprogramming prone condition (46 genes set; Fig. 4g). GO terms in green text are also induced by Neurog2 alone in EGF+bFGF culture. Abbreviations: FP, fluorescent protein; GO, Gene Ontology.

### Hmgb2-dependent expression of a specific set of neuronal maturation genes is necessary for efficient direct glia to neuron conversion

Our data suggested that low Hmgb2 expression levels in the bFGF culture could decrease astrocyte to neuron conversion via several non-mutually exclusive mechanisms: a) failure to activate the full neurogenic program induced in EGF+bFGF culture, b) prevention of the silencing of the conflicting alternative lineages and c) induction of a different neurogenic program from that in the EGF+bFGF culture. To directly test these possibilities, we analyzed the transcriptomic changes induced by the overexpression of Hmgb2 alone or in combination with Neurog2 in both bFGF and EGF+bFGF cultures.

Interestingly, Hmgb2 overexpression induced only several differentially expressed genes (DEGs) in either EGF+bFGF or bFGF cultures with respect to CAG-GFP control viral transduction ((Suppl. Fig. 6e, f; FC >2, padj < 0.05): two DEGs in the bFGF condition and four DEGs in the EGF+bFGF culture condition, Suppl. Table 2). This transcriptomic analysis, together with the lack of change in the conversion rate after Hmgb2 overexpression in both bFGF and EGF+bFGF astrocytes (Fig. 2d), suggested that Hmgb2 did not implement any specific neurogenic program on its own. Notably, the overexpression of Hmgb2 together with Neurog2 in the bFGF culture, as compared with control viral transduction, induced 255 genes (Fig. 3 g). This gene set was significantly enriched in GO biological processes associated with neural development, neuronal migration, axon guidance and synaptic signaling (Fig. 4f), similarly to the GO biological processes induced by Neurog2 alone in the EGF+bFGF condition (Fig. 4a). In addition, we observed downregulation of 164 genes (Suppl. Table 3) enriched in regulation of cell adhesion, actin filament organization, stress fiber assembly, and regulation of protein phosphorylation (Suppl. Fig. 6g), thus suggesting that simultaneous overexpression of Neurog2 and Hmgb2 suppresses gene expression that may block successful conversion of astroglia to neurons, possibly through post-translational modifications [57]. However, the down-regulated genes were not associated with specific glial or alternative fates induced by Neurog2 in the bFGF culture (Suppl. Fig. 6g).

To determine whether the dual overexpression of Neurog2+Hmgb2 might trigger similar transcriptional programs in the bFGF culture and the Neurog2-transduced the EGF+bFGF culture, we compared induced genes among three conditions: reprogramming prone culture (EGF+bFGF transduced with Neurog2 vs control virus), reprogramming resistant culture (bFGF transduced with Neurog2 vs control virus) and revived reprogramming culture (bFGF transduced with Neurog2+Hmgb2 vs control virus). We identified 88 genes that were shared across all three conditions (Fig. 4g) and were enriched in GO biological processes associated with neurogenesis, neuronal differentiation and migration, and trans-synaptic signaling (Suppl. Fig. 6h), in line with our findings that all conditions at least partially induced the neurogenic program. Furthermore, 46 genes (for example, *Prox1*, *Lrp8*, *Shf* and *Dscaml1*) were shared exclusively between the reprogramming prone conditions (bFGF Neurog2+Hmgb2 and EGF+bFGF Neurog2). This gene set was enriched in GO biological processes associated with axonogenesis, positive regulation of neurogenesis, neuron projection guidance, and nervous system development, thus implying that the upregulation of genes induced by the simultaneous overexpression of Neurog2 and Neurog2+Hmgb2 in the bFGF culture are associated with the acquisition of a more mature neuronal phenotype.

Together, our data suggested that the Hmgb2 protein aids in implementing the Neurog2-dependent, neurogenic program in astrocytes by facilitating the induction of a specific set of neurogenic, neuronal maturation-associated genes.

### Hmgb2 increases the chromatin accessibility of regions associated with the neurogenic program

We hypothesized that the establishment of the full neurogenic program by high levels of Hmgb2 is associated with Hmgb2-dependent chromatin changes. Therefore, we performed assay for transposase-accessible chromatin with high-throughput sequencing (ATAC-seq) on the cells from the same sorting samples used to generate transcriptomic libraries (Suppl. Fig. 5). We first examined the genome-wide chromatin accessibility profile at transcription start sites (TSSs ± 3.0 Kb) in both bFGF and EGF+bFGF cultures after the overexpression of Hmgb2, Neurog2, Neurog2+Hmgb2 and CAG-GFP control. The accessibility profile of Hmgb2 overexpressing astrocytes was comparable to that of the control regardless of the culture condition (Fig. 5a), in line with the lack of changes in the transcriptome and conversion rate analysis (Fig. 2e; Suppl. Fig. 6e, f). We did not observe any discernible increase in chromatin accessibility with simultaneous overexpression of Neurog2+Hmgb2 compared with Neurog2 in EGF+bFGF culture. However, we observed a substantial increase in chromatin accessibility after simultaneous overexpression of Neurog2+Hmgb2 compared with Neurog2 in the bFGF culture (Fig. 5b). This increase in TSS (±3 kb) accessibility might have been due to at least two mutually non-exclusive mechanisms: a) widespread TSS opening after Hmgb2 overexpression, or b) lineage specific changes. Therefore, we analyzed the TSS accessibility of neuronal cell-type-specific genes [58] (Fig. 5c). Whereas we observed the accessibility of these sites increased after both Neurog2 and Neurog2+Hmgb2 overexpression in the EGF+bFGF culture condition, in the bFGF culture condition, the increase in these sites was detectable only after simultaneous overexpression of Neurog2+Hmgb2 but not Neurog2 alone (Fig. 5c). Interestingly, the TSS opening was comparable between bFGF and EGF+bFGF astrocytes after Neurog2+Hmgb2 overexpression (Fig. 5c), in line with an increased conversion rate. Next, we wondered whether the Hmgb2-dependent increase in accessibility might be confined to neuronal genes or whether it might also occur in genes specific for other cell lineages. Therefore, we analyzed the dependence of the promoter accessibility of genes identifying ES cells [59,60], endothelial cells [61–63], and microglial cells [64,65] on Hmgb2 levels in bFGF culture (Fig. 5d). We found no significant differences in accessibility between the Hmgb2, Neurog2 or Neurog2+Hmgb2 treated astrocytes and the controls, thus indicating that the accessibility change after Neurog2+Hmgb2 overexpression was specific for neuronal fate.

**Figure 5.**
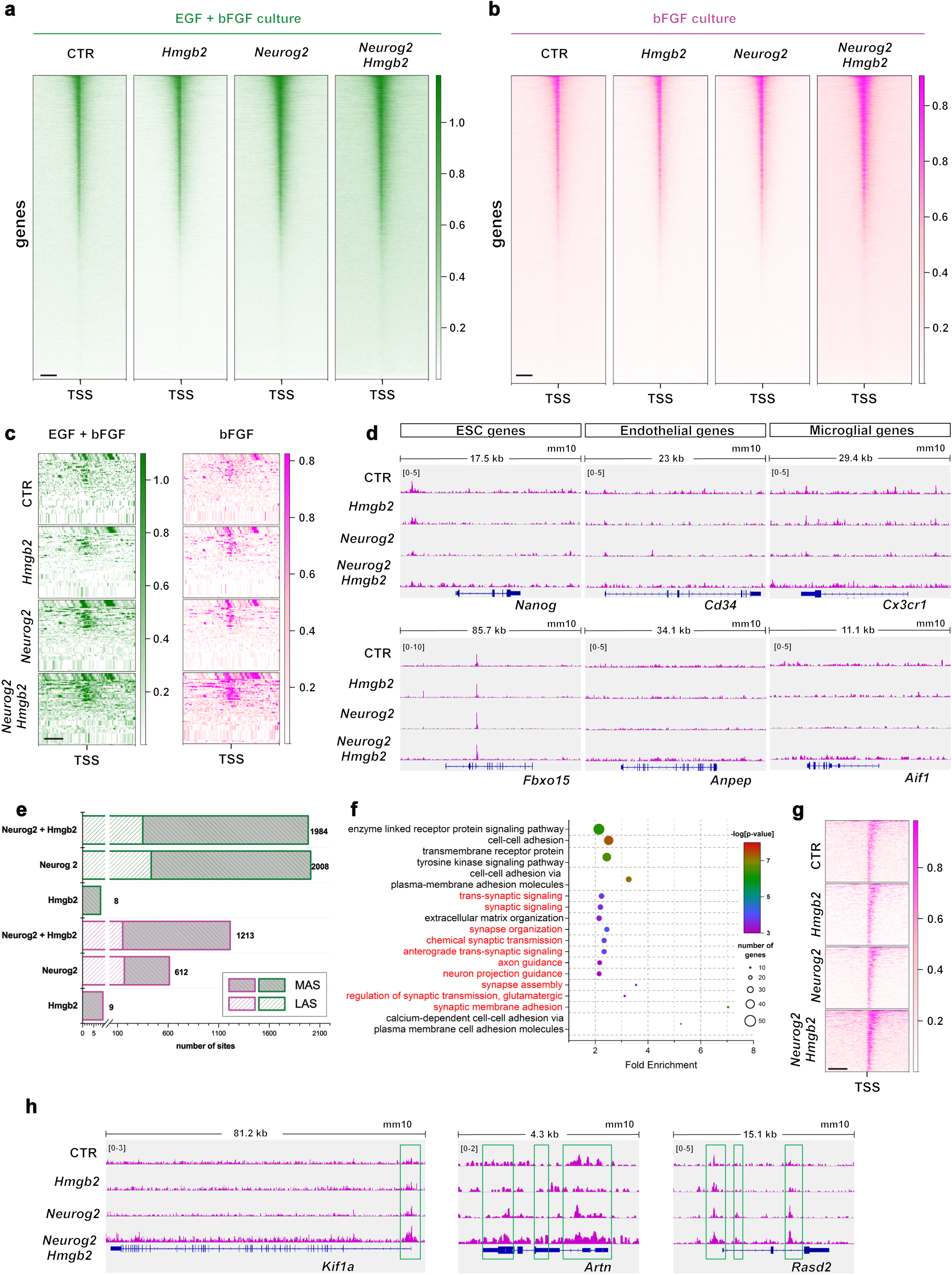
Hmgb2 improves the capability of Neurog2 to open promoters of neuronal maturation-associated genes. (**a,b**) Heat maps depicting opening of promoters by Neurog2 and Hmgb2 or their combination in EGF+bFGF (green, a) and bFGF (magenta, b) culture. Scale: 1 kb (**c**) Heat maps depicting ATAC signals in the promoters of the core neurogenic genes (Fig. 4d) 48 h after Neurog2, Hmgb2 or Neurog2+Hmgb2 overexpression in EGF+bFGF and bFGF cultures. (**d**) IGV tracks showing the ATAC signal in the promoters of genes identifying non-neuronal lineages 48 h after Neurog2, Hmgb2 or Neurog2+Hmgb2 overexpression in bFGF culture. (**e**) Histogram depicting the number of more (MAS) or less (LAS) accessible sites identified by ATAC 48 h after Neurog2, Hmgb2 or Neurog2+Hmgb2 overexpression in EGF+bFGF (green) and bFGF (magenta) cultures. (**f**) Plot depicting enriched GO biological process terms in the promoter set opened by Neurog2+Hmgb2 in bFGF culture 48 hours after viral transduction. (**g**) Heat map showing ATAC signal in the promoters of neuronal maturation related genes (red in panel e) 48 h after Neurog2, Hmgb2 or Neurog2+Hmgb2 overexpression in bFGF culture. (g) IGV tracks showing the ATAC signal in the promoters of representative genes involved in neuronal maturation 48 h after Neurog2, Hmgb2 or Neurog2+Hmgb2 overexpression in FGF culture. Green boxes indicate differentially accessible sites.

To identify direct conversion relevant changes in chromatin accessibility dependent on Hmgb2 levels, we determined the significant differentially accessible sites (DASs) after overexpression of Neurog2 and Neurog2+Hmgb2, compared with CAG-GFP-transduced cells, in the bFGF and EGF+bFGF culture conditions. In the bFGF culture, Neurog2 overexpression resulted in 612 DASs (445 more accessible sites (MASs) and 167 less accessible sites (LASs); Fig. 5e, Suppl. Table 4). Combined overexpression of Neurog2+Hmgb2 in the bFGF culture resulted in 1213 DASs (1062 MASs and 151 LASs; Fig. 5e, Suppl. Fig. 7a). However, this increase in accessibility did not change the accessibility profile induced by Neurog2 and Neurog2+Hmgb2 in the bFGF culture, because we observed a similar distribution of MAS in the gene bodies, promoters and intergenic regions (Suppl. Fig. 7b, c). Importantly, the Hmgb2-associated increase in MASs was not observed in EGF+bFGF astrocyte culture (Fig 5e), in agreement with our transcriptome analysis. To reveal the processes influenced by MASs, we analyzed genes associated with these sites (defined as genes within 3 kb upstream and downstream of the MAS) in GO analysis. MASs induced by the simultaneous overexpression of Neurog2+Hmgb2 in the bFGF culture were associated with nervous system development, synaptic membrane adhesion, axon guidance, synapse assembly and chemical synaptic transmission (Fig. 5f, Suppl. Table 5). This finding suggests that Hmgb2 (together with Neurog2) increases the accessibility of genes involved in neuronal maturation. Indeed, the promoters of synapse-associated genes such as *Kif1a* [66,67], *Artn* [68] and *Rasd2* [69] were closed in the bFGF culture after either control viral transduction or Hmgb2 overexpression (Fig. 5h), in line with the astrocytic fate of these cells. Moreover, Neurog2+Hmgb2 overexpression opened the synapse-associated promoters to a significantly greater extent than Neurog2 alone (Fig. 5g, h). We then asked whether the chromatin opening state of all or only a subset of Neurog2-induced maturation genes depended on the expression of Hmgb2. Therefore, we compared the MASs induced by Neurog2 in the two conversion prone conditions (overexpression of Neurog2 in EGF+bFGF and overexpression of Neurog2+Hmgb2 in bFGF culture) with MASs induced by Neurog2 in the conversion resistant condition (overexpression of Neurog2 in bFGF culture). We identified 395 MASs commonly induced in both conversion prone conditions (Fig. 6a). These MASs were enriched in processes associated with synapse formation (GO biological processes such as nervous system development, synaptic organization, trans-synaptic signals, potassium transport, and synaptic membrane adhesion; Fig. 6b, Suppl. Table 6). Importantly, the increase in the accessibility of these synapse-associated loci correlated with the increased expression of these genes after Neurog2+Hmgb2 overexpression in bFGF culture (Suppl. Fig. 8 a, b). However, we also observed 268 MASs induced by Neurog2 in all three conditions (Fig. 6a) that were enriched in synaptic processes (Fig. 6c, Suppl. Table 6). Therefore, these data suggested that the chromatin containing only a subset of genes associated with neuronal maturation was dependent on Hmgb2. However, the accessibility of these genes appeared to be instrumental for direct conversion.

**Figure 6.**
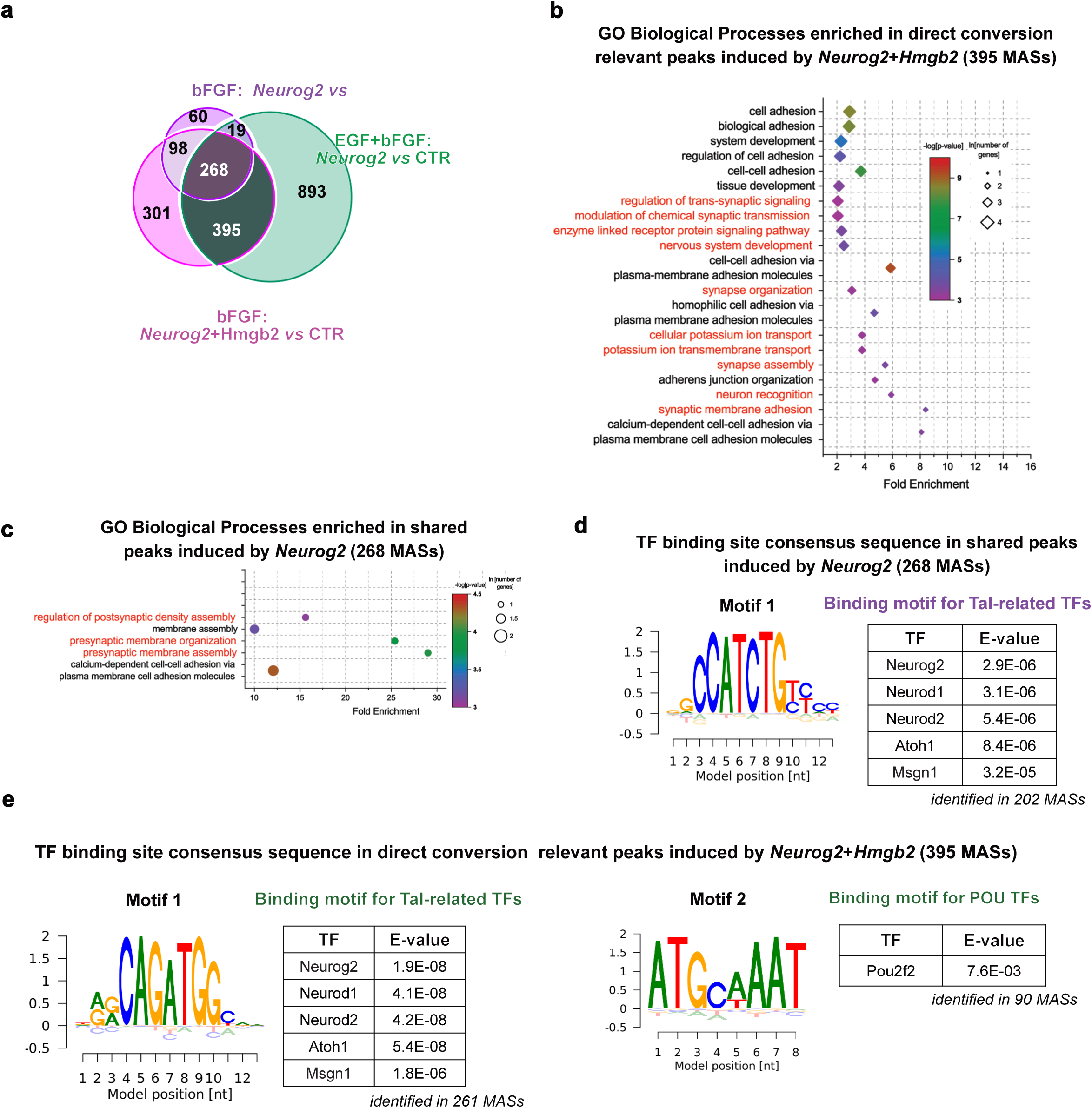
Hmgb2-dependent promoters contain an E-box and Pou2f2 factor binding motif. (**a**) Venn diagram illustrating the overlap in ATAC signals for MASs after Neurog2 overexpression in EGF+bFGF and bFGF cultures, with MASs induced by Neurog2 and Hmgb2 overexpression in bFGF culture 48 h after viral transduction. (**b**, **c**) Plots depicting enriched GO biological process terms in 395 peak set MASs in panel a (b) and 268 peak set MASs in panel a (b). (**d, e**) Transcription factor consensus sequences identified in 268 peak set MASs in panel a (d) and 395 peak set MASs in panel a (e), identified with *de novo* motif analysis. The motif image from the BaMM web server shows the likelihood of each nucleotide at each motif position. The color intensity reflects the probability, with darker colors indicating higher probabilities. Tables show transcription factors binding these motifs. Abbreviations: MAS, more accessible site; TF, transcription factor.

Together, our data supported a model in which Hmgb2 fosters the establishment of the full neurogenic program by increasing the accessibility and consequently the expression of neuronal maturation genes, thus leading to improved neuronal maturation.

### Hmgb2-dependent chromatin sites contain both E-boxes and Pou factor binding sites important for neuronal maturation

HMG proteins play a major role in controlling gene expression by increasing chromatin accessibility [70–72]. Therefore, we sought to identify the potential TF binding motifs enriched in the Hmgb2-dependent set of MASs (395 sites in Fig. 6a). To do so, we performed *de novo* motif enrichment analysis using BaMMmotif software. Motifs containing the consensus binding sequence of the Tal-associated TF family (Neurod1, Neurog2, Neurod2, Atoh1 and Msgn1) were enriched in Hmgb2-dependent set of MASs (Fig. 6d, Suppl. Table 7). In addition, we identified the motif that best matched the consensus sequence of the TF family of POU domain factors, such as Pou2f2 (Fig. 6e, Suppl. Table 7). Pou2f2 is a direct Neurog2 target [73] and has been reported to be involved in the implementation of proper neuronal identity [74,75]. This finding suggested that in the bFGF culture, some of the E-box motif sites bound by Neurog2 (Tal related factors) were inaccessible, but with the addition of Hmgb2, these sites became accessible, thereby increasing Neurog2-binding and enhancing reprogramming efficiency. Additionally, we investigated MASs with consensus binding sequences for both Tal-associated factors (Neurog2) and POU domain factors. We identified that 56 of 395 MASs contained binding motifs for both TF families, and were associated with neuronal maturation (GO processes: regulating actin filaments assembly, chemotaxis, and potassium ion transport; Suppl. Fig. 8d and Suppl. Table 7), including the Robo-Slit pathway. Robo-Slit pathway has been reported to regulate not only axonal pathfinding but also neuronal maturation [76]. Moreover, we observed enrichment in genes associated with the negative regulation of proliferation, thus possibly improving the terminal differentiation of converted cells. Interestingly, *de novo* motif analysis of the common 268 Neurog2-induced MASs identified the binding motif of the TF family of Tal-associated factors, but not of the POU domain factors (Fig. 6d). These data suggested that Hmgb2 levels set the lineage barrier by controlling the accessibility of both the direct Neurog2 targets and targets of TFs downstream of Neurog2, such as Pou3f2 or Neurod.

To directly test the importance of Hmgb2 in neuronal maturation, we analyzed the neurite complexity of the converted neurons in the conversion prone cultures (overexpression of Neurog2 in EGF+bFGF and overexpression of Neurog2+Hmgb2 in bFGF culture) and the conversion resistant culture (overexpression of Neurog2 in bFGF culture) in induced neurons with Sholl analysis 7 days after viral transduction (Fig. 7a). Indeed, Neurog2-induced neurons in the bFGF culture showed fewer intersections than the Neurog2-induced neurons in the EGF+bFGF culture (Fig. 7b, c). Lower neurite complexity is indicative of less mature neurons. The complexity of neurites in neurons generated from bFGF astrocytes by the combined overexpression of Neurog2 and Hmgb2 increased compared to overexpression of Neurog2 only. These converted neurons were indistinguishable from those generated by overexpression of Neurog2 in the EGF+bFGF-cultured astrocytes (Fig. 7b, c).

**Figure 7.**
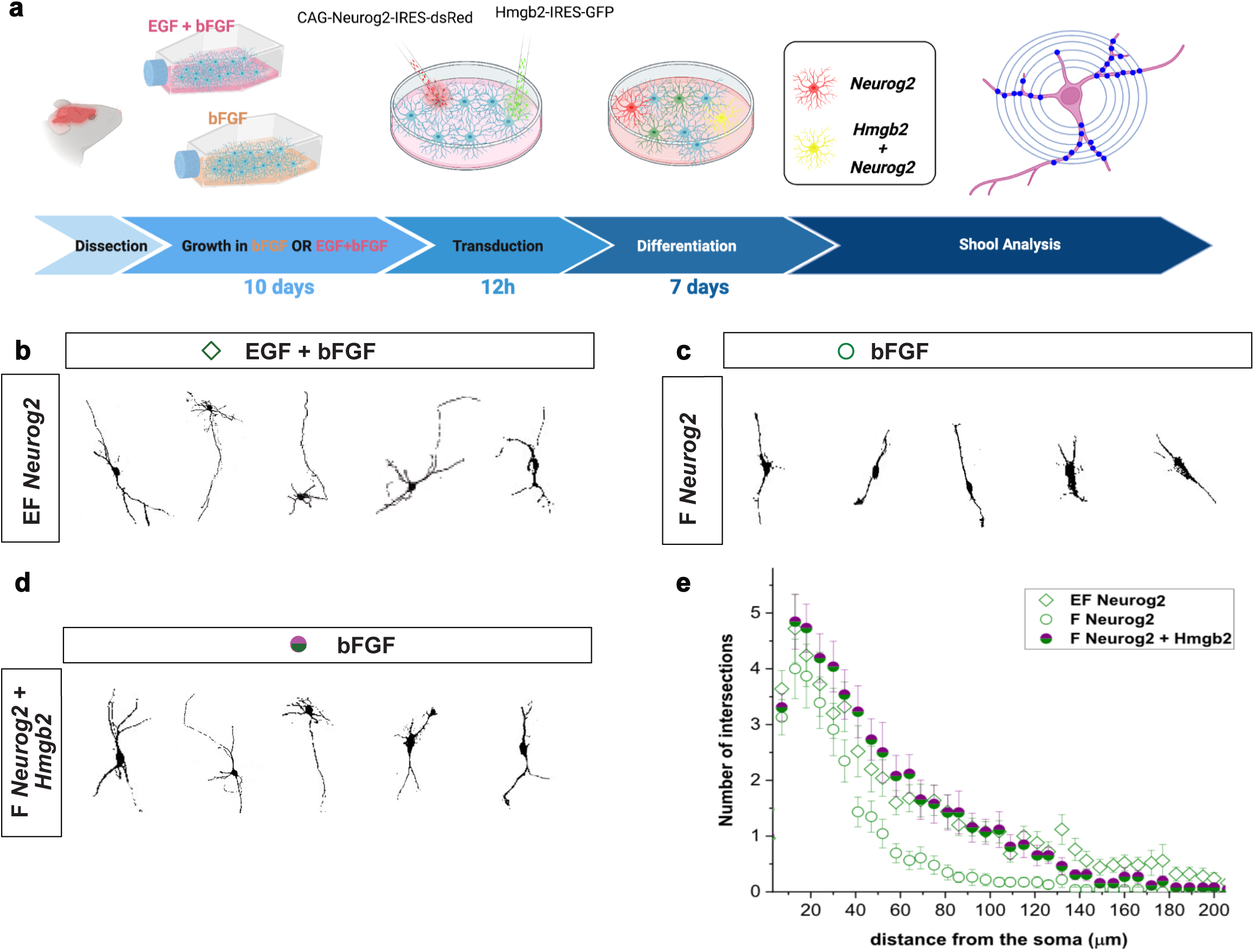
Hmgb2 and Neurog2 overexpression increases complexity of iN. (**a**) Scheme depicting the experimental paradigm used for Shool analysis. (**b**) Representative thresholded images of neuronal cells used for Sholl analysis. (**c**) Sholl analysis of induced neurons by concurrent overexpression of Neurog2 and Neurog2+Hmgb2 in EGF and EGF+bFGF culture 7 days after viral transduction. Abbreviations: MAS, more accessible site; TF, transcription factor.

## Discussion

The establishment of neuronal identity during direct astrocyte to neuron conversion is achieved in very different environmental context from that of the bona fide neurogenesis occurring during embryonic development or in adult brain neurogenic niches [49,51]. This includes not only the different starting populations [49] but also the unique signaling milieus [77–79]. The growth factors released after injury regulate the conversion process, including neuronal maturation and neural circuit repair. Here, we presented a novel *in vitro* system to study the influence of growth factors on fate conversion. Using this system, we showed that EGF, potentially provided by the injured environment, is necessary for efficient neuronal conversion and proper maturation via the regulation of the chromatin binding protein Hmgb2. In combination with several different neurogenic fate determinants, Hmgb2 is capable of inducing the full neurogenic program, as indicated by Hmgb2 gain and loss of function experiments. Our model predicted that prolonged injury-induced elevation in bFGF levels decreased the reprogrammability of astrocytes to neurons. However, the FGF signal per se did not prevent the induction of a set of processes associated with neurogenesis and neuronal fate in astrocytes during Neurog2-mediated conversion. This finding is in line with reports that the FGF promotes neurogenesis [80–82], although the neuronal subtypes generated in such context differ [82]. Importantly, the chromatin states in direct conversion and during embryonic neurogenesis may differ: the chromatin states during neurogenesis require fewer re-arrangements in embryonic development, because large numbers of neurogenic gene loci in radial glial cells, the neuronal stem cells of the developing CNS, are already in an open configuration [83,84]. Interestingly, genes involved in synapse formation and neuronal maturation are already in an active chromatin state without detectable gene expression in both radial glia and committed neuronal progenitors [83,85], thus implying the existence of an active inhibitory mechanism keeping the progenitor state primed toward neurogenesis and preventing their premature differentiation. Importantly, Hmgb2 opens the loci of these classes of genes during astrocyte to neuron conversion, thus supporting the concept that overexpression of Neurog2+Hmgb2 endows postnatal astrocytes with some stem cell features. This concept is also in line with the expression of Hmgb2 during activation of quiescent neural stem cells in the adult brain [53] and its role in adult neurogenesis [86]. However, we did observe immediate expression of synaptic genes in postnatal astrocytes without the maintenance of these primed neuronal states, thus suggesting that the mechanisms preventing premature differentiation operating in the neuronal stem cells are not established during astrocyte to neuron conversion. This possibility reinforces the concept that direct neuronal conversion does not fully recapitulate the developmental trajectory underlying neuronal differentiation [44,48]. Instead, the overexpression of reprogramming factors induces early re-arrangements of chromatin along with changes in gene expression. However, during late morphological and functional maturation stages of the induced neurons, changes in chromatin are negligible [87]. Moreover, in our *in vitro* system, we did not observe any changes in astrocyte proliferation due to the overexpression of Hmgb2 alone or in combination with different neurogenic TFs, thus further limiting the spectrum of neural stem cell features induced in the postnatal astrocytes. Interestingly, Hmgb2 induces similar chromatin changes in postnatal astrocytes to the HMG group protein A2, a different HMG-box-containing family member in gliogenic radial glial cells. These chromatin changes are sufficient to prolong the neurogenic phase during cortical development and lead to the generation of new postnatal neurons [88]. During this period, progenitors normally generate glial cells, thus potentially implicating similar mechanisms in the Hmga2-mediated extension of neurogenic period and the Hmgb2-mediated direct astrocyte to neuron conversion. Because Hmga2 is associated with Polycomb signaling [89], testing whether the same system would be operational during the Hmgb2-dependent conversion should prove interesting, because Ezh2 maintains the lineage barriers during fibroblast to neuron conversion [90]. Both Hmgb2 and Hmga2 bind AT-rich DNA segments with little to no sequence specificity [91][71]. Nevertheless, we observed highly specific Hmgb2-dependent opening of chromatin containing late neuronal maturation genes, thus prompting questions regarding HMG protein binding specificity. This specificity could be provided by an interacting protein, e.g., neurogenic TF Neurog2, because we observed an enrichment of the typical E-box binding sequence in the promoters when Hmgb2 was overexpressed in astrocytes. However, our findings did not reveal a direct interaction of Hmgb2 with Neurog2 via WB or mass spectrometry, thus making this scenario unlikely. An alternative explanation may be that Hmgb2 stabilizes the regulatory loops (transactivation domains, TADs) involved in the expression of synaptic genes. The regulatory roles of such domains have been demonstrated for neurogenesis downstream of Neurog2 during embryonal cerebral cortex development [92]. Moreover, both Hmgb2 and Hmga2 have been implicated in TAD establishment [93,94]. The stabilization of regulatory loops induced by Neurog2 may indeed provide a mechanistic explanation for the Hmgb2-dependent opening of chromatin regions containing the Neurog2 binding E-boxes. These data further challenge the common belief that Neurog2 is a pioneer TF. In contrast to the on-target pioneering function of Ascl1 during reprogramming [87,95], in fibroblast to neuron conversion, Neurog2 requires additional factors, such as forskolin and dorsomorphin or Sox4, that are necessary for not only late neuronal maturation but also the induction of early reprogramming changes [73,96]. We demonstrated that, at least in the case of astrocyte to neuron conversion, Neurog2 function is dependent on Hmgb2. Because Hmgb2 increases the accessibility of various sites, including the binding motif of the Neurog2 target Pou2f2 [92], our data suggested that Neurog2 must open the chromatin of maturation genes that are transcriptionally regulated by direct Neurog2 targets. Our study provides mechanistic insights into previously described improvements in neuronal reprogramming with the infusion of EGF and FGF [37]. Interestingly, EGF and FGF exhibit different temporal dynamics post-injury, with a very narrow expression window and a presumably diminished activity window of EGF [41]. This window correlates with the expression of Hmgb2, thus suggesting that prolonged expression the either EGF or Hmgb2 after TBI might be important in the success of neuronal replacement therapies. Furthermore, our model may also explain the lower direct conversion rates induced by Neurog2 in some starting cellular populations, such as oligodendrocyte precursor cells [97], in which the promoters might not yet be open. Similarly, such multilevel control is compatible with the ability of Neurog2 to induce different neuronal subtypes or maturation stages in different, permissive starting cells [46,96,98,99], given that maturation loci defining the neuronal subtype could be differentially accessible for Neurog2 direct targets.

Interestingly, the overexpression of Neurog2 in bFGF-grown astrocytes induced not only a partial neurogenic program but also additional transcriptional programs associated with alternative fates, such as cartilage formation and immune cell differentiation. The induction of alternative fates or a failure to repress the original fate can lead to abortive conversion and concomitant death of reprogrammed cells [100], thereby possibly mechanistically explaining the lower Neurog2-mediated conversion efficiency in the bFGF culture. Because Hmgb2 overexpression does not specifically repress the astrocytic fate, yet significantly improves the conversion efficiency, the abortive direct conversion is unlikely to explain the lower efficiency in direct conversion. Interestingly, we did not observe Hmgb2-dependent opening of regions associated with alternative fate genes, thus supporting the idea that alternative fate induction is independent of the Hmgb2-induced changes in chromatin states. Hmgb2-dependent changes in the transcription rate [101], RNA stability or RNA splicing could account for the enrichment of alternative fates observed in mRNA analysis, because Hmgb2 has been proposed to have an RNA-binding domain [91]. Importantly, we observed changes in chromatin opening for only genes associated with the neurogenic lineage.

## Conclusions

Together, our results provide a mechanistic framework for translating environmental signals into a specific program involved in neuronal maturation downstream of the neurogenic fate determinants via chromatin modification. Interestingly, this aspect of neuronal reprogramming is the least understood and stands to be further improved, particularly *in vivo*.

## Suppl. Figure Legends

**Suppl. Figure 1. Growth conditions define the direct conversion rate.**

**(a)** Scheme depicting the experimental paradigm used for astrocyte to neuron conversion. (**b-e**) Micrographs depicting the fate of transduced cells after control viral transduction in EGF+bFGF (b), bFGF (c), EGF (d) culture and Neurog2 overexpression in EGF culture (e) 7 days after viral transduction. Scale bars: 50 µm. (**f**, **g**) Dot plots showing direct conversion efficacy of Neurog2 overexpression in EGF culture (f) as well as Pou2f2, and Pou3f2+Hmgb2 overexpression in EGF+bFGF and bFGF culture (g). Data are shown as median±IQR; single dots represent independent biological replicates. Significance was tested with two-tailed Mann-Whitney test. Abbreviations: FP, fluorescent protein.

**Suppl. Figure 2: Characterization of the starting population in EGF+bFGF and bFGF culture.**

**(a)** Scheme depicting the experimental paradigm used to characterize initially transduced cells. (**b**, **c**, **d**, **e**, **g**, **h**, **j**, **k**, **l**, **m**) Micrographs illustrating identity assessment of control virally transduced cells 24 h after transduction. Yellow arrows indicate identity marker positive transduced, GFP-positive cells. Scale bars: 50 µm. (**f, i, n**) Dot plots showing the proportion of transduced cells with the indicated identity. Data are shown as median±IQR; single dots represent independent biological replicates. Significance was tested with two-tailed Mann-Whitney test.

**Suppl. Figure 3: The growth factor induced barrier is reversible.**

**(a)** Scheme depicting the experimental paradigm used to address the stability of the growth factor induced lineage barrier. (**b**-**e**) Micrographs illustrating the identity of control virus (b, d) and Neurog2-encoding virus (c, e) transduced cells cultured first in bFGF and then in EGF+bFGF (b, c), and of cells cultured first in EGF+bFGF and then bFGF (d, e). Identity assessment was performed 7 days after viral transduction. Scale bar in b-e: 50 μm. (**f**) Dot plots showing the proportions of transduced cells acquiring neuronal identity 7 days after viral transduction. Data are shown as median±IQR; single dots represent independent biological replicates. Significance was tested with two-tailed Mann-Whitney test.

**Suppl. Figure 4: Traumatic brain injury induces Hmgb2 expression in gray matter reactive astrocytes.**

**(a)** Scheme depicting the experimental paradigm. (**b**-**c’**) Micrographs showing the expression of Hmgb2 in the intact (b) and injured hemisphere (c) 5 days after injury. (c’) Orthogonal projections of the optical Z-stack depicting the expression of Hmgb2 in astrocytes of the injured hemisphere. Scale bars in b and c 100 μm and in c’ 10 μm.

**Suppl. Figure 5: Isolation of transduced cells for RNAseq and ATACseq.**

**(a)** Scheme depicting the workflow used to isolate transduced cells 48 h after transduction for omic analysis. (**b**) Plots demonstrating the FACS sorting gates and settings used to sort cells transduced with control, Neurog2 and Hmgb2 expressing viruses.

**Suppl. Figure 6. Neurog2+Hmgb2 overexpression in bFGF culture induces a transcriptional subset necessary for successful direct conversion.**

(**a-b**) Volcano plots of differentially expressed genes (DEGs) induced by Neurog2 in EGF+bFGF culture (a) and bFGF culture (b) 48 hours after viral transduction. (**c**) Plot depicting enriched GO biological processes of 96 shared genes (Fig. 4c) induced by Neurog2 in both EGF+bFGF and bFGF culture 48 hours after viral transduction. (**d**) Plot depicting enriched GO biological processes of uniquely induced genes by Neurog2 in EGF+bFGF culture (225 gene set; in Fig. 4c, green text) and bFGF culture (41 gene set in Fig. 4c, magenta text) 48 hours after viral transduction. (**e**, **f**) Volcano plot of DEGs induced by Hmgb2 in EGF+bFGF culture (f) and bFGF culture (g) 48 hours after viral transduction. (**g**) Plot depicting enriched GO biological processes of genes downregulated by Neurog2+Hmgb2 overexpression in bFGF culture 48 hours after viral transduction. Red text highlights processes associated with cytoskeletal remodeling, and blue depicts processes involved in adhesion. (**h**) Plot depicting enriched GO biological processes of the gene set commonly induced by Neurog2 in EGF+bFGF, bFGF culture and by Neurog2+Hmgb2 in bFGF culture (88 genes in Fig. 4g). Black text highlights processes associated with neurogenesis.

**Suppl. Figure 7. Hmgb2 increases the ability of Neurog2 to open chromatin in bFGF culture.**

**(a)** Heat map depicting accessibility of MASs induced by Hmgb2 (9 MASs), Neurog2 (445 MASs) and the combination of Neurog2+Hmgb2 (1062 MASs) in bFGF culture 48 h after viral transduction. Scale: 1 kb. (**b-c**) Pie charts of genomic distribution of MASs induced by Neurog2 (b) and the combination of Neurog2+Hmgb2 (c) in bFGF culture 48 h after viral transduction.

**Suppl. Figure 8. Additional sites opened by Hmgb2 and Neurog2 overexpression are associated with the establishment of synaptic contacts and/or maturation of neurons.**

**(a)** IGV tracks showing the ATAC signals of genes associated with synapse formation/function 48 h after viral transduction in bFGF culture. Boxes indicate signals significantly broadened by co-expression of Neurog2 and Hmgb2. (**b**) Box plots depicting expression of synapse-associated genes (from panel a) after control, Neurog2, Hmgb2 and Neurog2+Hmgb2 overexpression in bFGF culture 48 hours after viral transduction. (**c**) Venn diagram illustrating the overlap of MASs with the Tal-associated factor binding motif (motif 1, E-box) and POU domain factor binding motif (motif 2, POU) induced by Neurog2 in EGF+bFGF culture and induced by Neurog2+Hmgb2 in bFGF culture. (**d**) Plot depicting GO biological processes enriched in genes with promoters containing binding motifs for both Tal-associated factors and POU domain factors (56 promoters in c).

## Suppl. Table Legends

**Suppl. Table 1. GO analysis of processes enriched in the EGF+bFGF and bFGF only proteomes.**

**Suppl. Table 2. Full list of differentially regulated genes between different conditions.**

**Suppl. Table 3. GO analysis associated with RNA-seq analysis.**

**Suppl. Table 4. Full list of MAS and DAS with their genomic location.**

**Suppl. Table 5. GO analysis associated with ATAC analysis.**

**Suppl. Table 6. GO analysis associated ATAC peaks enriched in different reprogramming conditions.**

**Suppl. Table 7. Full list of MAS and DAS with Neurog2 and Pou TF binding motifs.**

## Material and Methods

### Experimental animals

Experiments were conducted on both, female and male animals, which were either wild types (C57BL/6J mice) or transgenic Hmgb2−/− animals on a C57BL/6 background [102]. The Hmgb2−/− mice do not show gross phenotypical abnormalities and do not differ to wild-type siblings (Ronfani et al., 2001). For all *in vitro* experiments, animals at postnatal stage P5-P6 were used. Injuries were done in adult 8-10 weeks old animals. Animals were kept under standard conditions with access to water and food ad libitum. All animal experimental procedures were performed in accordance with the German and European Union guidelines and were approved by the Institutional Animal Care and Use Committee (IACUC) and the Government of Upper Bavaria under license number: AZ 55.2-1-54-2532-171-2011 and AZ 55.2-1-54-2532-150-11. All efforts were made to minimize animal suffering and to reduce the number of animals used.

### Stab wound injury

Prior to every surgery, mice were deeply anesthetised by intra-peritoneal injection of sleep solution (Medetomidin (0,5mg/kg) / Midazolam (5mg/kg) / Fentanyl (0,05mg/kg)) complemented by local lidocaine application (20 mg/g). After the injection of the anaesthesia, mice were checked for pain reactions by pinching their tail and toes. Stab wound injury was performed in the somatosensory cortex, as previously described [97,103]. The following coordinates relative to Bregma were used: medio-lateral: 1,0 μm; rostro-caudal: −1,2 μm to - 2,2 μm; dorso-ventral: −0,6 μm. Anaesthesia was antagonized with an subcutan injection of awake solution (Atipamezol (2,5mg/kg) / Flumazenil (0,5mg/kg) / Buprenorphin (0,1mg/kg)) and the mice were kept on a pre-warmed pad until they were awake and recovered from the surgery.

### Perfusion and tissue section preparation

Prior to perfusion, animals were deeply anesthetized with overdoses of cocktail of ketamine (100 mg/kg) / xylazine (10 mg/kg). Subsequently, they were transcardially perfused first with cold PBS, followed by fresh ice-cold 4% PFA in PBS for 20 minutes. The brain was then removed from the skull, post-fixed in the same fixative overnight at 4 °C, cryoprotected in 30% sucrose and cut at the cryostat at 40 μm tick sections.

### Preparation of PDL-coated glass coverslips

Glass coverslips were washed first with acetone and boiled for 30 min in ethanol containing 0,7% (v/v) HCl. After two washing steps with 100% ethanol, coverslips were dried at RT and autoclaved for 2 h at 180 °C. Coverslips were washed with D-PBS and coated with poly-D-lysine (PDL, 0.02 mg/ml) solution for at least 2 h at 37 °C. Following coating, coverslips were washed three times with autoclaved ultrapure water, dried in the laminar flow and stored at 4 °C until needed.

### Primary culture of postnatal cortical astroglial cells

Postnatal cortical astroglia were isolated and cultured as described previously [104]. Following decapitation of postnatal (P5-P6) wild-type C57BL/6J mice, the skin and the skull were removed, and the brain was extracted avoiding any tissue damage and placed into the 10 mM HEPES solution for dissection. After separating the two hemispheres, the meninges was removed and white matter of cerebral cortex was dissected using fine forceps and collected in a tube with astrocyte medium (Fetal calf serum-FCS (10% (v/v)); Horse serum-HS (5% (v/v)); glucose (3,5 mM); B27 supplement; Penicillin/Streptomycin (100 I.U/ml Pen and 100 μg/ml Strep) in DMEM/F12+GlutaMAX). The tissue was mechanical dissociated with a 5 ml pipette and placed into uncoated plastic flasks for cell expansion in astrocyte medium supplemented with the two growth factors EGF (10 ng/ml) + bFGF (10 ng/ml each) or with bFGF (10 ng/ml) only as specified for each experiment. After 4-5 days, the medium was exchanged and supplied with the fresh growth factors. After 10 days of culturing, cultured cells were rinsed with DPBS and contaminating oligodendrocyte precursor cells were removed by brusquely shaking the culture flasks several times. Astroglial cells were then detached from the flask by trypsinization and seeded onto poly-D-lysine (PDL)-coated glass coverslips at a density of 8×10^4^ cells per well in a 24-well plate with astrocyte medium for immunohistochemical analysis. For the ATAC-seq and RNA-seq experiments, cells were plated in T75 flasks with a seeding density of 3×10^6^ cells per flask. 2-4 h after seeding, the cells were transduced with different retroviral vectors in a ratio of 1 μl virus per 1 ml medium to prevent virus toxicity. Astrocyte medium was changed 12-18 h after viral transduction to differentiation medium (glucose (3,5 mM); B27 supplement; Penicillin/Streptomycin (100 I.U/ml Pen and 100 μg/ml Strep) in DMEM/F12+GlutaMAX) containing neither EGF nor bFGF up to the immunocytochemical analysis timepoint. The cells were cultured as indicated in each experiment. Cells were fixed in cold 4% PFA for 20 min and rinsed with cold D-PBS before immunocytochemical analysis.

For the ATAC-seq and RNA-seq experiments, the cells were kept in the astrocyte medium and collected 48 h after viral transduction. Astrocytes were detached from the flask by trypsinization, prepared for the FACS and sorted for the following ATAC-seq and RNA-seq experiments according to the fluorophore expression. The astroglial cultures from the Hmgb2−/− transgenic animals were prepared as described above, however, the cortical tissue from each animal was kept separately and placed into the small T25 flask. In addition, the tips of the tails were used for genotyping as described in [102]. The cultures from Hmgb2−/− transgenic mice were grown only in the double growth factor condition containing EGF+bFGF.

### Immunocytochemistry and immunohistochemistry

Immunostaining was performed on cell culture samples or free-floating brain sections. Specimens were treated with blocking buffer (0,5% Triton-X-100; 10% normal goat serum (NGS) in D-PBS) to reduce non-specific binding. The same buffer was used to dilute the primary antibodies. The specimens were incubated with the primary antibody mixture overnight at 4°C (brain tissue) °C or for 2 hours at RT (cell culture samples), followed by 3x 10 min washing steps with PBS. In order to visualize primary antibody binding, samples were exposed to appropriate species and/or subclass specific secondary antibodies conjugated to Alexa Fluor 488, 546 or 647 (Invitrogen) for about 90 min at RT protected from light. Secondary antibodies were diluted 1:1000 in blocking buffer. Nuclei were visualized with DAPI (4’,6-diamidino-2-phenylindole) that was added to the mix of secondary antibodies. Following extensive washing steps with PBS, coverslips or sections were mounted with Aqua Poly/Mount (Polysciences) and imaged.

Following primary antibodies were used: Chick-anti-GFP (Aves Lab, GFP-120; 1:1000); Rabbit-anti-RFP (Rockland, 600-401-379; 1:500); Mouse IgG1-anti-GFAP (Sigma-Aldrich, G3893; 1:500); Rabbit-anti-GFAP (DakoCytomation, Z0334; 1:1000); Mouse IgG1κ-anti-S100β (Sigma-Aldrich, S2644; 1:500); Rabbit-anti-OLIG2 (Thermo Fischer, AB9610; 1:500); Mouse IgG2a-anti-αSMA (Sigma-Aldrich, A2547; 1:400); Rabbit-anti-Ki67 (Abcam, 15580; 1:200); Rat-anti-Ki67 (DakoCytomation, M7249; 1:200); Rabbit-anti-PH3 (Ser10) (Thermo Fischer, 06-570; 1:200); Guinea pig-anti-DCX (Thermo Fischer, AB-2253; 1:1000); Mouse IgG2b-anti-β-III-TUBULIN (Sigma-Aldrich, T8660; 1:500); Mouse IgG1-anti-NEUN (Chemicon, MAB 377; 1:250); Rabbit-anti-HMGB2 (Abcam, ab67282; 1:1000); Mouse IgG2aκ-anti-HMGB2 (Sigma-Aldrich, 07173-3E5; 1:500); Mouse IgG2aκ-anti-HMGB2 antibody requires termal (15 min at 95°C) antigen retrieval using the citrate buffer (10 mM; pH 6). Primary antibody binding was revealed using class-specific secondary antibody coupled to Alexa fluorophore (Invitrogen, Germany). All secondary antibodies were used at dilution 1:1000.

### Image acquisition and quantifications

Immunostainings were analysed with a fluorescent Microscope Axio Imager M2m (Zeiss) using the ZEN software (Zeiss) with a 20x or 40x objective. Fluorescent-labelled sections were photographed with FV1000 confocal laser-scanning microscope (Olympus), using the FW10-ASW 4.0 software (Olympus). The quantifications of *in vitro* cultured cells were performed using the ZEN software (Zeiss) analysing at least 25 randomly taken pictures per coverslip depending on the number of transduced cells. In total, 100-200 retroviral vector-transduced cells were quantified from randomly chosen fields on a single coverslip. 3 coverslips in each experiment (biological replicate) were analysed. The number of experiments is indicated in corresponding Figure. The number of induced neurons was expressed as a percentage out of all transduced cells.

To analyse the number of apoptotic cells, between 350-550 DAPI labelled cells were counted from 5 randomly selected fields on one coverslip.

In the reprogramming experiments of the astrocytes isolated from Hmgb2+/+, Hmgb2+/−- and Hmgb2−/− animals, each of the single animals was considered as a biological replicate and at least 3 coverslips were counted per animal. We analysed in total 6 litters containing wild-type, heterozygous or homozygous littermates.

Western blots using the Fiji software as previously described [105]. All lanes of interest were outlined using the rectangular selection tool and the signal intensity of each band was calculated by determining the area under the peak. The measurements of the corresponding α-ACTIN bands were used to normalize the amount of proteins loaded on the gel.

### Sholl Analysis

We analysed only DCX positive cells 7 days after viral transduction. Single cells were isolated and subjected to Sholl analysis using the ImageJ plug-in ‘Sholl Analysis’. We used the following parameters: starting radius 5μm; ending radius 500 μm; radius step size 5 μm. The number of crossings per cell were visualized and analysed using Origin.

### FACS analysis and sorting

Astrocytes were collected by trypsinization 48 h after retroviral transduction, washed, resuspended in DPBS and analysed using a FACS Aria II instrument (BD Biosciences) in the FACSFlowTM medium. Debris and aggregated cells were gated out by forward-scatter area (FSC-A) and side-scatter area (SSC-A). Forward scatter area (FSC-A) vs. forward scatter width (FSC-W) was used to discriminate doublets from single cells. To set the gates for the sorting, untransduced astrocytes were recorded. Sorted cells were collected in DPBS, counted and divided into two batches: 50000 cells were immediately processed for ATAC-seq and the remaining cells were collected for RNA-seq library preparation.

### ATAC-sequencing

Assay for Transposase Accessible Chromatin with high-throughput sequencing (ATAC-seq), a method to detect genome-wide chromatin accessibility, was performed following the published protocol [106,107]. Briefly, right after the FACS sorting, 50000 cells were lysed, the nuclei were extracted and resuspended with the transposase reaction mix (25 μl 2x TD buffer (Illumina); 2,5 μl Transposase (Illumina); 22,5 μl nuclease free water), following by transposition reaction for 30 minutes at 37°C °C. To stop the transposition reaction, samples were purified using a Qiagen MinElute PCR (Qiagen) purification kit according to the manufacturer instructions. Open chromatin fragments were first amplified for 5 cycles and then for additional 7-8 cycles, as determined by RT-qPCR, using the combination of primer Ad1_noMX (5’ AATGATACGGCGACCACCGAGATCTACACTCGTCGGCAGCGTCAGATGTG 3’) and the Nextera Index Kit (Illumina) primer N701-N706. Libraries were purified using a Qiagen MinElute PCR purification kit (Qiagen) and their quality was assessed using the Bioanalyzer High-Sensitivity DNA kit (Agilent) according to the manufacturer’s instructions. The concentration of each library was measured by Qubit using the provided protocol. Libraries were pooled for sequencing and the pool contained 20 ng of each library. Prior to sequencing, pooled libraries were additionally purified with AMPure beads (ratio 1:1) to remove contaminating primer dimers and quantified using Qubit and the Bioanalyzer High-Sensitivity DNA kit (Agilent). 50-bp paired-end deep sequencing was carried out on HiSeq 4000 (Illumina).

### ATAC-sequencing analysis

For the analysis of bulk ATAC-seq data, we followed the Harvard FAS Informatics ATAC-seq guidelines. The quality of raw FASTQ reads were checked using FastQC (Version 0.11.9). The low quality read (< 20bp) and adapter sequences were trimmed by Cutadapt (Version 4.0). The trimmed reads were mapped to the mouse reference genome (mm10) by using Bowtie2 (parameter: --very-sensitive -X 1000 --dovetail). Samtools were then used to convert and sort the sam files into bam files. Peak calling step was performed with Genrich for each sample separately to identify accessible regions. Genrich peak caller has a mode (-j) assigned to ATAC-Seq analysis mode and allows running all of the post-alignment steps via peak-calling with one command. Mitochondrial reads and PCR duplicates were removed by -e chrM and -r argument respectively. To generate count table matrix for differential analysis bam2counts (intePareto R-based package) was used to count reads fall into specific genomic positions by importing all the bam files and merging all the bed files into one (importing GenomicRanges and GenomicAlignments libraries). DESeq2 (version 1.26.0) was used for differential accessibility analysis of the count data. The relatively more open and closed sites are called MAS and LAS respectively (fold change (FC) > 2 and adjusted P-value < 0.05) and the annotation of these sites were performed using R-based packages Chip-seeker (TSS ± 3.0 Kb) (version 1.28.3). For visualization, the bamcovage deeptools (version 3.5.1) were used to normalize the data by importing the scaling factor from DESeq2 (version 1.36.0). The normalized bigwig files used to visualize the coverage using deeptools and samtools. These bigwig files were loaded into the IGV tool to visualize the peak at the gene level. The Venn diagrams were made using the BioVenn web application tool. The Gorilla tool was used to generate the GO Biological processes, with a cut-off of enrichment > 2 and p-value of < 0.01.

### Motif analysis

BaMMmotif (https://bammmotif.soedinglab.org/home/) was used to perform *de novo* motif enrichment analysis by providing MASs fasta sequence [108] as input and all detected accessible sites fasta sequences as background using default parameters. We selected the motifs with an AvRec score above 0.5 as candidates for further analysis. The mouse database HOCOMOCO v11 was used for motif annotation, and the most significant transcription factors matching the motif with e-values below 0.001 were considered as potential binders.

### Preparation of libraries for RNA-sequencing

Sorted cells were resuspended in 100 μl extraction buffer of the PicoPureTM RNA isolation kit (Thermo Fischer Scientific) and the RNA was extracted according to the manufacturer’s instructions. The Agilent 2100 Bioanalyzer was used to assess RNA quality and concentration. For the RNA-seq library preparation, only high-quality RNA with RIN values >8 were used. cDNA was synthesized from 10 ng of total RNA using SMART-Seq v4 Ultra Low Input RNA Kit (Takara Bio), according to the manufacturer’s instructions. The total number of amplification cycles was determined by RT-qPCR side reaction according to manufacturer’s instruction. PCR-amplified cDNA was purified by immobilization on AMPure XP beads. Prior to generating the final library for sequencing, the Covaris AFA system was used to perform cDNA shearing in Covaris microtubes (microTUBE AFA Fiber Pre-Slit Snap-Cap 6×16mm), resulting in 200-500 bp long cDNA fragments that were subsequently purified by ethanol precipitation. Prior to library preparation using the MicroPlex Library Preparation kit v2 (Diagenode) according to the user manual, the quality and concentration of the sheared cDNA were assessed using an Agilent 2100 Bioanalyzer. Final libraries were evaluated using an Agilent 2100 Bioanalyzer and the concentration was measured with Qubit Fluorometer (Thermo Fischer Scientific). The uniquely barcoded libraries were multiplexed onto one lane and 100-bp paired-end deep sequencing was carried out at the HiSeq 4000 (Illumina) generating ∼20 million reads per sample.

### Transcriptome data analysis (Bulk RNA Seq)

The raw paired-end FASTQ files were mapped to the mouse reference genome (mm10) using STAR RNA-seq aligner (version 2.7.2b). Aligned reads in the BAM files were then quantified by HTSeq-count (Version 0.9.1) based on annotation file GENCODE Release M25 (GRCm38.p6). The gene-level count matrix was imported into the R/Bioconductor package DESeq2 (version 1.26.0) for normalization and differential expression with FC > 2, adjusted P-value < 0.05. Venn diagrams were created using the web application BioVenn tool and heatmaps were generated using gplots and RColorBrewer R-based/Bioconductor tools. For GO enrichment analysis of the assigned set of genes we used the GOrilla tool by providing background genes. The enriched GO term (biological processes) possessing enrichment > 2, containing at least 1% of the input genes and p-value specified in the figure legend were visualized using Origin.

### Protein isolation and Western blot

Postnatal cortical astroglia were isolated and cultured as described above. After 10 days of culturing with growth factors EGF+bFGF or bFGF, cells were detached from the flask by trypsinization, washed and counted. 0,5×10^6^ cells were lysed in RIPA buffer containing cOmplete Protease Inhibitor cocktail (Roche). Protein extraction and Western blotting is performed as previously described [109]. The following antibodies were used: Rabbit-anti-HMGB2 (Abcam, ab67282; 1:5000); Mouse-anti-ACTIN (Millipore, MAB1501; 1:10000); HRP-coupled anti-mouse IgG1 (GE Healthcare, NA931; 1:20000) and HRP-coupled anti-rabbit IgG (Jackson ImmunoResearch,111-036-045; 1:20000).

### Quantitative mass spectrometry

Treated adherent astrocytes were lysed and subjected to tryptic protein digest using a modified FASP protocol [110]. Proteomic measurements were performed on a LTQ Orbitrap XL mass spectrometer (Thermo Scientific) online coupled to an Ultimate 3000 nano-HPLC (Dionex). Peptides were enriched on a nano trap column (100 μm i.d. × 2 cm, packed with Acclaim PepMap100 C18, 5 μm, 100 Å, Dionex) prior to separation on an analytical C18 PepMap column (75 μm i.d. × 25 cm, Acclaim PepMap100 C18, 3 μm, 100Å, Dionex) in a 135 min linear acetonitrile gradient from 3% to 34% ACN. From the high resolution orbitrap MS pre-scan (scan range 300 – 1500 m/z), the ten most intense peptide ions of charge ≥ +2 were selected for fragment analysis in the linear ion trap if they exceeded an intensity of at least 200 counts. The normalized collision energy for CID was set to a value of 35. Every ion selected for fragmentation was excluded for 30 s by dynamic exclusion. The individual raw-files were loaded to the Progenesis software (version 4.1, Waters) for label free quantification and analyzed as described [111,112]. MS/MS spectra were exported as Mascot generic file and used for peptide identification with Mascot (version 2.4, Matrix Science Inc., Boston, MA, USA) in the Ensembl Mouse protein database (release 75, 51765 sequences). Search parameters used were as follows: 10 ppm peptide mass tolerance and 0.6 Da fragment mass tolerance, one missed cleavage allowed, carbamidomethylation was set as fixed modification, methionine oxidation and asparagine or glutamine deamidation were allowed as variable modifications. A Mascot-integrated decoy database search was included. Peptide assignments were filtered for an ion score cut-off of 30 and a significance threshold of p < 0.01 and were reimported into the Progenesis software. After summing up the abundances of all peptides allocated to each protein, resulting normalized protein abundances were used for calculation of fold-changes and corresponding p-values.

### Expression plasmids

In order to overexpress different neurogenic transcription factors in the astroglial cells, we used Moloney murine leukemia virus (MMLV)-derived retroviral vectors, expressing neurogenic fate determinants under the regulatory control of a strong and silencing-resistant pCAG promoter. All our construct encode a neurogenic factor followed by an internal ribosomal entry site (IRES) and either GFP or dsRED as reporter proteins, allowing simultaneous reporter expression. For control experiments, we used a retrovirus encoding for the fluorescent proteins (GFP or dsRED) behind the IRES driven by the same CAG promoter. We used the following expression vectors: pCAG-IRES-GFP [43]; pCAG-IRES-dsRED [43]; pCAG-Neurog2-IRES-dsRED [43]; pCAG-Pou3f2 -IRES-dsRED [113]; pCAG-Sox11-IRES-GFP [46]; pCAG-Hmgb2-IRES-GFP^(this^ ^work)^.

### Cloning pCAG-Hmgb2-IRES-GFP construct

cDNA for Hmgb2 were synthetized at Genscript, containing BamHI and HindIII in order to clone them into the pENTR1A entry vector. The cDNAs were then transferred to the retroviral destination vector pCAG-IRES-dsRED/GFP using the Gateway cloning method (Invitrogen) according to the manufacturer’s instructions. The correct sequence was confirmed using Sanger sequencing before viral vector production.

### Retroviral vector production

The VSV-G-pseudotyped retroviruses were prepared using the HEK293-derived retroviral packaging cell line (293GPG) (Ory et al., 1996) that stably express the gag-pol genes of murine leukemia virus and vsv-g under the control of a tet/VP16 transactivator as previously described (Heinrich et al., 2011). The viral particles were stored in TNE (Tris-HCl pH=7,8 (50mM); NaCl (130mM); EDTA (1mM)) buffer at −80 0C until use.

### Statistical analysis

Numbers of biological replicates can be seen on the dot plots or in the figure legend in case of the bar charts. All results are presented as median ± interquartile range (IQR). IQR was calculated in RStudio [114], using the default method based on type 7 continuous sample quantile. For the reprogramming experiments, statistical analysis was performed in Origin using non-parametric Mann-Whitney U test unless differently specified for particular experiments.

## Declarations

### Availability of data and materials

Proteome data set is available at PRIDE database (https://www.ebi.ac.uk/pride/). The dataset identifier is PXD044288. During the review process the data could be accessed using the following username: and password: C9naS7jL.

The RNAseq and ATACseq datasets are available at Gene Expression Omnibus (GEO). The accession number is pending. The reviewer token will be provided upon request.

### Authors’ contributions

P.M., T.L. and J. N. conceived the project and designed experiments. A.S.-M., V.S., F.B., and J.N. performed experiments. J. M.-P. and S.M.H. analyzed proteome. L.R. and M. B. provided Hmgb2 KO animals. P. M. and J.N. wrote the manuscript with input from all authors.

## Acknowledgments

We thank all members of the Neurogenesis and Regeneration group for experimental input, discussions and critical reading of the manuscript. We acknowledge the support of the following core facilities: the Bioimaging Core Facility at the BioMedical Center of LMU Munich and the Sequencing Facility at the Helmholtz Zentrum München.

## References

1. Barker RA, Götz M, Parmar M. New approaches for brain repair—from rescue to reprogramming. Nature 2018 557:7705 [Internet]. 2018 [cited 2022 Aug 16];557:329–34. Available from: https://www.nature.com/articles/s41586-018-0087-1

2. Götz M, Bocchi R. Neuronal replacement: Concepts, achievements, and call for caution. Curr Opin Neurobiol [Internet]. 2021 [cited 2022 Aug 18];69:185–92. Available from: https://pubmed.ncbi.nlm.nih.gov/33984604/

3. Torper O, Götz M. Brain repair from intrinsic cell sources: Turning reactive glia into neurons. Prog Brain Res. 2017.

4. Griesbach GS, Masel BE, Helvie RE, Ashley MJ. The Impact of Traumatic Brain Injury on Later Life: Effects on Normal Aging and Neurodegenerative Diseases. J Neurotrauma. 2018;35.

5. Bocchi R, Masserdotti G, Götz M. Direct neuronal reprogramming: Fast forward from new concepts toward therapeutic approaches. Neuron [Internet]. 2022 [cited 2022 Aug 18];110:366–93. Available from: https://pubmed.ncbi.nlm.nih.gov/34921778/

6. Zhou Y, Shao A, Yao Y, Tu S, Deng Y, Zhang J. Dual roles of astrocytes in plasticity and reconstruction after traumatic brain injury. Cell Communication and Signaling. BioMed Central Ltd.; 2020.

7. Amamoto R, Arlotta P. Development-inspired reprogramming of the mammalian central nervous system. Science (1979) [Internet]. 2014;343:1239882. Available from: http://www.ncbi.nlm.nih.gov/pubmed/24482482

8. Berninger B, Costa MR, Koch U, Schroeder T, Sutor B, Grothe B, et al. Functional properties of neurons derived from in vitro reprogrammed postnatal astroglia. J Neurosci [Internet]. 2007;27:8654–64. Available from: http://www.ncbi.nlm.nih.gov/entrez/query.fcgi?cmd=;Retrieve&db=;PubMed&dopt=;Citation&list_uids=;17687043

9. Buffo A, Vosko MR, Erturk D, Hamann GF, Jucker M, Rowitch D, et al. Expression pattern of the transcription factor Olig2 in response to brain injuries: implications for neuronal repair. Proc Natl Acad Sci U S A [Internet]. 2005/12/07. 2005;102:18183–8. Available from: http://www.ncbi.nlm.nih.gov/entrez/query.fcgi?cmd=Retrieve&db=PubMed&dopt=Citation&list_uids=16330768

10. Grande A, Sumiyoshi K, López-Juárez A, Howard J, Sakthivel B, Aronow B, et al. Environmental impact on direct neuronal reprogramming in vivo in the adult brain. Nat Commun [Internet]. 2013;4. Available from: 10.1038/ncomms3373

11. Heinrich C, Blum R, Gascón S, Masserdotti G, Tripathi P, Sánchez R, et al. Directing Astroglia from the Cerebral Cortex into Subtype Specific Functional Neurons. McKay RDG, editor. PLoS Biol [Internet]. 2010/05/27. 2010;8:e1000373. Available from: http://www.ncbi.nlm.nih.gov/entrez/query.fcgi?cmd=Retrieve&db=PubMed&dopt=Citation&list_uids=20502524

12. Herrero-Navarro Á, Puche-Aroca L, Moreno-Juan V, Sempere-Ferràndez A, Espinosa A, Susín R, et al. Astrocytes and neurons share region-specific transcriptional signatures that confer regional identity to neuronal reprogramming. Sci Adv. 2021;7.

13. Liu F, Zhang Y, Chen F, Yuan J, Li S, Han S, et al. Neurog2 directly converts astrocytes into functional neurons in midbrain and spinal cord. Cell Death Dis. 2021;12.

14. Masserdotti G, Gillotin S, Sutor B, Drechsel D, Irmler M, Jørgensen HF, et al. Transcriptional Mechanisms of Proneural Factors and REST in Regulating Neuronal Reprogramming of Astrocytes. Cell Stem Cell [Internet]. 2015;17:74–88. Available from: https://linkinghub.elsevier.com/retrieve/pii/S1934590915002234

15. Mattugini N, Bocchi R, Scheuss V, Russo GL, Torper O, Lao CL, et al. Inducing Different Neuronal Subtypes from Astrocytes in the Injured Mouse Cerebral Cortex. Neuron [Internet]. 2019;103:1086–1095.e5. Available from: https://linkinghub.elsevier.com/retrieve/pii/S0896627319306932

16. Ninkovic J, Steiner-Mezzadri A, Jawerka M, Akinci U, Masserdotti G, Petricca S, et al. The BAF complex interacts with Pax6 in adult neural progenitors to establish a neurogenic cross-regulatory transcriptional network. Cell Stem Cell. 2013;13.

17. Niu W, Zang T, Zou Y, Fang S, Smith DK, Bachoo R, et al. In vivo reprogramming of astrocytes to neuroblasts in the adult brain. Nat Cell Biol [Internet]. 2013;15:1164–75. Available from: http://www.ncbi.nlm.nih.gov/pubmed/24056302

18. Pereira M, Birtele M, Shrigley S, Benitez JA, Hedlund E, Parmar M, et al. Direct Reprogramming of Resident NG2 Glia into Neurons with Properties of Fast-Spiking Parvalbumin-Containing Interneurons. Stem Cell Reports [Internet]. 2017;9:742–51. Available from: https://www.ncbi.nlm.nih.gov/pubmed/28844658

19. Zhang L, Lei Z, Guo Z, Pei Z, Chen Y, Zhang F, et al. Development of Neuroregenerative Gene Therapy to Reverse Glial Scar Tissue Back to Neuron-Enriched Tissue. Front Cell Neurosci. 2020;14.

20. Vierbuchen T, Wernig M. Direct lineage conversions: unnatural but useful? Nat Biotechnol [Internet]. 2011/10/15. 2011;29:892–907. Available from: http://www.ncbi.nlm.nih.gov/entrez/query.fcgi?cmd=Retrieve&db=PubMed&dopt=Citation&list_uids=21997635

21. Lujan E, Chanda S, Ahlenius H, Sudhof TC, Wernig M. Direct conversion of mouse fibroblasts to self-renewing, tripotent neural precursor cells. Proc Natl Acad Sci U S A [Internet]. 2012/02/07. 2012;109:2527–32. Available from: http://www.ncbi.nlm.nih.gov/entrez/query.fcgi?cmd=Retrieve&db=PubMed&dopt=Citation&list_uids=22308465

22. Marro S, Pang ZP, Yang N, Tsai MC, Qu K, Chang HY, et al. Direct lineage conversion of terminally differentiated hepatocytes to functional neurons. Cell Stem Cell [Internet]. 2011/10/04. 2011;9:374–82. Available from: http://www.ncbi.nlm.nih.gov/entrez/query.fcgi?cmd=Retrieve&db=PubMed&dopt=Citation&list_uids=21962918

23. Treutlein B, Lee QY, Camp JG, Mall M, Koh W, Shariati SA, et al. Dissecting direct reprogramming from fibroblast to neuron using single-cell RNA-seq. Nature [Internet]. 2016;534:391–5. Available from: https://www.ncbi.nlm.nih.gov/pubmed/27281220

24. Pfisterer U, Kirkeby A, Torper O, Wood J, Nelander J, Dufour A, et al. Direct conversion of human fibroblasts to dopaminergic neurons. Proceedings of the National Academy of Sciences [Internet]. 2011;108:10343–8. Available from: http://www.ncbi.nlm.nih.gov/pubmed/21646515

25. Wapinski OL, Vierbuchen T, Qu K, Lee QY, Chanda S, Fuentes DR, et al. Hierarchical mechanisms for direct reprogramming of fibroblasts to neurons. Cell [Internet]. 2013;155:621–35. Available from: http://www.ncbi.nlm.nih.gov/pubmed/24243019

26. Wapinski OL, Lee QY, Chen AC, Li R, Corces MR, Ang CE, et al. Rapid Chromatin Switch in the Direct Reprogramming of Fibroblasts to Neurons. Cell Rep [Internet]. 2017;20:3236–47. Available from: https://www.ncbi.nlm.nih.gov/pubmed/28954238

27. Noack F, Vangelisti S, Raffl G, Carido M, Diwakar J, Chong F, et al. Multimodal profiling of the transcriptional regulatory landscape of the developing mouse cortex identifies Neurog2 as a key epigenome remodeler. Nat Neurosci. 2022;25.

28. Pataskar A, Jung J, Smialowski P, Noack F, Calegari F, Straub T, et al. NeuroD1 reprograms chromatin and transcription factor landscapes to induce the neuronal program. Embo J [Internet]. 2016;35:24–45. Available from: https://www.ncbi.nlm.nih.gov/pubmed/26516211

29. Heinrich C, Bergami M, Gascón S, Lepier A, Dimou L, Sutor B, et al. Sox2-mediated conversion of NG2 glia into induced neurons in the injured adult cerebral cortex. Stem Cell Reports [Internet]. 2014;in press:1000–14. Available from: http://www.ncbi.nlm.nih.gov/pubmed/25458895

30. Liu M-L, Zang T, Zou Y, Chang JC, Gibson JR, Huber KM, et al. Small molecules enable neurogenin 2 to efficiently convert human fibroblasts into cholinergic neurons. Nat Commun [Internet]. 2013;4. Available from: 10.1038/ncomms3183

31. Russo GL, Sonsalla G, Natarajan P, Breunig CT, Bulli G, Merl-Pham J, et al. CRISPR-Mediated Induction of Neuron-Enriched Mitochondrial Proteins Boosts Direct Glia-to-Neuron Conversion. Cell Stem Cell. 2021;28.

32. Gascón S, Murenu E, Masserdotti G, Ortega F, Russo GL, Petrik D, et al. Identification and Successful Negotiation of a Metabolic Checkpoint in Direct Neuronal Reprogramming. Cell Stem Cell [Internet]. 2016;18:396–409. Available from: https://linkinghub.elsevier.com/retrieve/pii/S1934590915005482

33. Hu X, Qin S, Huang X, Yuan Y, Tan Z, Gu Y, et al. Region-Restrict Astrocytes Exhibit Heterogeneous Susceptibility to Neuronal Reprogramming. Stem Cell Reports. 2019;12.

34. Addington CP, Roussas A, Dutta D, Stabenfeldt SE. Endogenous Repair Signaling after Brain Injury and Complementary Bioengineering Approaches to Enhance Neural Regeneration: Supplementary Issue: Stem Cell Biology. Biomark Insights. 2015.

35. Sun D, Bullock MR, McGinn MJ, Zhou Z, Altememi N, Hagood S, et al. Basic fibroblast growth factor-enhanced neurogenesis contributes to cognitive recovery in rats following traumatic brain injury. Exp Neurol. 2009;216.

36. Ninomiya M, Yamashita T, Araki N, Okano H, Sawamoto K. Enhanced neurogenesis in the ischemic striatum following EGF-induced expansion of transit-amplifying cells in the subventricular zone. Neurosci Lett. 2006;403.

37. Grande A, Sumiyoshi K, López-Juárez A, Howard J, Sakthivel B, Aronow B, et al. Environmental impact on direct neuronal reprogramming in vivo in the adult brain. Nat Commun [Internet]. 2013;4. Available from: 10.1038/ncomms3373

38. Hung LY, Tseng JT, Lee YC, Xia W, Wang YN, Wu ML, et al. Nuclear epidermal growth factor receptor (EGFR) interacts with signal transducer and activator of transcription 5 (STAT5) in activating Aurora-A gene expression. Nucleic Acids Res. 2008;36.

39. Choi HS, Choi BY, Cho YY, Mizuno H, Kang BS, Bode AM, et al. Phosphorylation of histone H3 at serine 10 is indispensable for neoplastic cell transformation. Cancer Res. 2005;65.

40. Patel NS, Rhinn M, Semprich CI, Halley PA, Dollé P, Bickmore WA, et al. FGF Signalling Regulates Chromatin Organisation during Neural Differentiation via Mechanisms that Can Be Uncoupled from Transcription. PLoS Genet. 2013;9.

41. Addington CP, Roussas A, Dutta D, Stabenfeldt SE. Endogenous Repair Signaling after Brain Injury and Complementary Bioengineering Approaches to Enhance Neural Regeneration: Supplementary Issue: Stem Cell Biology. Biomark Insights. 2015.

42. Heinrich C, Blum R, Gascon S, Masserdotti G, Tripathi P, Sanchez R, et al. Directing astroglia from the cerebral cortex into subtype specific functional neurons. PLoS Biol [Internet]. 2010/05/27. 2010;8:e1000373. Available from: http://www.ncbi.nlm.nih.gov/entrez/query.fcgi?cmd=Retrieve&db=PubMed&dopt=Citation&list_uids=20502524

43. Heinrich C, Gotz M, Berninger B. Reprogramming of postnatal astroglia of the mouse neocortex into functional, synapse-forming neurons. Methods Mol Biol [Internet]. 2012;814:485–98. Available from: http://www.ncbi.nlm.nih.gov/pubmed/22144327

44. Gascon S, Masserdotti G, Russo GL, Gotz M. Direct Neuronal Reprogramming: Achievements, Hurdles, and New Roads to Success. Cell Stem Cell [Internet]. 2017;21:18–34. Available from: http://www.ncbi.nlm.nih.gov/pubmed/28686866

45. Hack MA, Sugimori M, Lundberg C, Nakafuku M, Götz M, Gotz M. Regionalization and fate specification in neurospheres: the role of Olig2 and Pax6. Mol Cell Neurosci [Internet]. 2004/04/15. 2004;25:664–78. Available from: http://www.ncbi.nlm.nih.gov/pubmed/15080895

46. Masserdotti G, Gillotin S, Sutor B, Drechsel D, Irmler M, Jørgensen HF, et al. Transcriptional Mechanisms of Proneural Factors and REST in Regulating Neuronal Reprogramming of Astrocytes. Cell Stem Cell. 2015;

47. Russo GL, Sonsalla G, Natarajan P, Breunig CT, Bulli G, Merl-Pham J, et al. CRISPR-Mediated Induction of Neuron-Enriched Mitochondrial Proteins Boosts Direct Glia-to-Neuron Conversion. Cell Stem Cell. 2021;28.

48. Masserdotti G, Götz M. A decade of questions about the fluidity of cell identity. Nature. 2020;578.

49. Bocchi R, Masserdotti G, Götz M. Direct neuronal reprogramming: Fast forward from new concepts toward therapeutic approaches. Neuron. 2022.

50. Gascón S, Masserdotti G, Russo GL, Götz M. Direct Neuronal Reprogramming: Achievements, Hurdles, and New Roads to Success. Cell Stem Cell. 2017.

51. Ninkovic J, Götz M. Fate specification in the adult brain - lessons for eliciting neurogenesis from glial cells. BioEssays. 2013;35.

52. Ninkovic J, Götz M. Understanding direct neuronal reprogramming — from pioneer factors to 3D chromatin. Curr Opin Genet Dev. 2018;52.

53. Kimura A, Matsuda T, Sakai A, Murao N, Nakashima K. HMGB2 expression is associated with transition from a quiescent to an activated state of adult neural stem cells. Developmental Dynamics. 2018;247.

54. Mall M, Kareta MS, Chanda S, Ahlenius H, Perotti N, Zhou B, et al. Myt1l safeguards neuronal identity by actively repressing many non-neuronal fates. Nature [Internet]. 2017;544:245–9. Available from: https://www.ncbi.nlm.nih.gov/pubmed/28379941

55. Hsieh CY, Nakamura PA, Luk SO, Miko IJ, Henkemeyer M, Cramer KS. Ephrin-B reverse signaling is required for formation of strictly contralateral auditory brainstem pathways. Journal of Neuroscience. 2010;30.

56. Kania A, Klein R. Mechanisms of ephrin-Eph signalling in development, physiology and disease. Nat Rev Mol Cell Biol. 2016.

57. Hindley C, Ali F, McDowell G, Cheng K, Jones A, Guillemot F, et al. Post-translational modification of Ngn2 differentially affects transcription of distinct targets to regulate the balance between progenitor maintenance and differentiation. Development. 2012;139.

58. Zhang ZH, Jhaveri DJ, Marshall VM, Bauer DC, Edson J, Narayanan RK, et al. A comparative study of techniques for differential expression analysis on RNA-seq data. PLoS One. 2014;9.

59. Pan G, Thomson JA. Nanog and transcriptional networks in embryonic stem cell pluripotency. Cell Res. 2007.

60. Tokuzawa Y, Kaiho E, Maruyama M, Takahashi K, Mitsui K, Maeda M, et al. Fbx15 Is a Novel Target of Oct3/4 but Is Dispensable for Embryonic Stem Cell Self-Renewal and Mouse Development. Mol Cell Biol. 2003;23.

61. Petrovic N, Schacke W, Gahagan JR, O’Conor CA, Winnicka B, Conway RE, et al. CD13/APN regulates endothelial invasion and filopodia formation. Blood. 2007;110.

62. Mina-Osorio P, Winnicka B, O’Conor C, Grant CL, Vogel LK, Rodriguez-Pinto D, et al. CD13 is a novel mediator of monocytic/endothelial cell adhesion. J Leukoc Biol. 2008;84.

63. Lin G, Finger E, Gutierrez-Ramos JC. Expression of CD34 in endothelial cells, hematopoietic progenitors and nervous cells in fetal and adult mouse tissues. Eur J Immunol. 1995;25.

64. Jung S, Aliberti J, Graemmel P, Sunshine MJ, Kreutzberg GW, Sher A, et al. Analysis of Fractalkine Receptor CX 3 CR1 Function by Targeted Deletion and Green Fluorescent Protein Reporter Gene Insertion. Mol Cell Biol. 2000;20.

65. Schwab JM, Frei E, Klusman I, Schnell L, Schwab ME, Schluesener HJ. AIF-1 expression defines a proliferating and alert microglial/macrophage phenotype following spinal cord injury in rats. J Neuroimmunol. 2001;119.

66. Okada Y, Yamazaki H, Sekine-Aizawa Y, Hirokawa N. The neuron-specific kinesin superfamily protein KIF1A is a uniqye monomeric motor for anterograde axonal transport of synaptic vesicle precursors. Cell. 1995;81.

67. Niwa S, Tanaka Y, Hirokawa N. KIF1Bβ- and KIF1A-mediated axonal transport of presynaptic regulator Rab3 occurs in a GTP-dependent manner through DENN/MADD. Nat Cell Biol. 2008;10.

68. Wang R, Rossomando A, Sah DWY, Ossipov MH, King T, Porreca F. Artemin induced functional recovery and reinnervation after partial nerve injury. Pain. 2014;155.

69. Errico F, Santini E, Migliarini S, Borgkvist A, Centonze D, Nasti V, et al. The GTP-binding protein Rhes modulates dopamine signalling in striatal medium spiny neurons. Molecular and Cellular Neuroscience. 2008;37.

70. Bianchi ME, Agresti A. HMG proteins: Dynamic players in gene regulation and differentiation. Curr Opin Genet Dev. 2005.

71. Štros M. HMGB proteins: Interactions with DNA and chromatin. Biochim Biophys Acta Gene Regul Mech. 2010.

72. Thomas JO, Travers AA. HMG1 and 2, and related “architectural” DNA-binding proteins. Trends Biochem Sci. 2001.

73. Smith DK, Yang J, Liu ML, Zhang CL. Small Molecules Modulate Chromatin Accessibility to Promote NEUROG2-Mediated Fibroblast-to-Neuron Reprogramming. Stem Cell Reports. 2016;7.

74. Javed A, Mattar P, Lu S, Kruczek K, Kloc M, Gonzalez-Cordero A, et al. Pou2f1 and Pou2f2 cooperate to control the timing of cone photoreceptor production in the developing mouse retina. Development (Cambridge). 2020;147.

75. Harris A, Masgutova G, Collin A, Toch M, Hidalgo-Figueroa M, Jacob B, et al. Onecut factors and Pou2f2 regulate the distribution of V2 interneurons in the mouse developing spinal cord. Front Cell Neurosci. 2019;13.

76. Gonda Y, Namba T, Hanashima C. Beyond Axon Guidance: Roles of Slit-Robo Signaling in Neocortical Formation. Front Cell Dev Biol. 2020.

77. Hevner RF. From Radial Glia to Pyramidal-Projection Neuron: Transcription Factor Cascades in Cerebral Cortex Development. Mol Neurobiol [Internet]. 2006 [cited 2019 Nov 3];33:033–50. Available from: http://www.ncbi.nlm.nih.gov/pubmed/16388109

78. Sofroniew M V. Astrocyte Reactivity: Subtypes, States, and Functions in CNS Innate Immunity. Trends Immunol. 2020.

79. Burda JE, Sofroniew M V. Reactive gliosis and the multicellular response to CNS damage and disease. Neuron [Internet]. 2014;81:229–48. Available from: http://www.ncbi.nlm.nih.gov/pubmed/24462092

80. Kang W, Hébert JM. FGF signaling is necessary for neurogenesis in young mice and sufficient to reverse its decline in old mice. Journal of Neuroscience. 2015;35.

81. Goldshmit Y, Tang JKKY, Siegel AL, Nguyen PD, Kaslin J, Currie PD, et al. Different Fgfs have distinct roles in regulating neurogenesis after spinal cord injury in zebrafish. Neural Dev. 2018;13.

82. Smith KM, Fagel DM, Stevens HE, Rabenstein RL, Maragnoli ME, Ohkubo Y, et al. Deficiency in Inhibitory Cortical Interneurons Associates with Hyperactivity in Fibroblast Growth Factor Receptor 1 Mutant Mice. Biol Psychiatry. 2008;63.

83. Pataskar A, Jung J, Smialowski P, Noack F, Calegari F, Straub T, et al. NeuroD1 reprograms chromatin and transcription factor landscapes to induce the neuronal program. Embo J [Internet]. 2016;35:24–45. Available from: https://www.ncbi.nlm.nih.gov/pubmed/26516211

84. Noack F, Pataskar A, Schneider M, Buchholz F, Tiwari VK, Calegari F. Assessment and site-specific manipulation of DNA (hydroxy-)methylation during mouse corticogenesis. Life Sci Alliance. 2019;2.

85. Aprea J, Prenninger S, Dori M, Ghosh T, Monasor LS, Wessendorf E, et al. Transcriptome sequencing during mouse brain development identifies long non-coding RNAs functionally involved in neurogenic commitment. EMBO Journal. 2013;32.

86. Abraham AB, Bronstein R, Reddy AS, Maletic-Savatic M, Aguirre A, Tsirka SE. Aberrant Neural Stem Cell Proliferation and Increased Adult Neurogenesis in Mice Lacking Chromatin Protein HMGB2. PLoS One. 2013;8:e84838.

87. Wapinski OL, Lee QY, Chen AC, Li R, Corces MR, Ang CE, et al. Rapid Chromatin Switch in the Direct Reprogramming of Fibroblasts to Neurons. Cell Rep [Internet]. 2017;20:3236–47. Available from: https://www.ncbi.nlm.nih.gov/pubmed/28954238

88. Kishi Y, Fujii Y, Hirabayashi Y, Gotoh Y. HMGA regulates the global chromatin state and neurogenic potential in neocortical precursor cells. Nat Neurosci [Internet]. 2012;15:1127–33. Available from: http://www.ncbi.nlm.nih.gov/pubmed/22797695

89. Zhou X, Zhong S, Peng H, Liu J, Ding W, Sun L, et al. Cellular and molecular properties of neural progenitors in the developing mammalian hypothalamus. Nat Commun. 2020;11.

90. Lee SW, Oh YM, Lu YL, Kim WK, Yoo AS. MicroRNAs Overcome Cell Fate Barrier by Reducing EZH2-Controlled REST Stability during Neuronal Conversion of Human Adult Fibroblasts. Dev Cell. 2018;46.

91. Starkova T, Polyanichko A, Tomilin AN, Chikhirzhina E. Structure and Functions of HMGB2 Protein. Int J Mol Sci. 2023;24:8334.

92. Noack F, Vangelisti S, Raffl G, Carido M, Diwakar J, Chong F, et al. Multimodal profiling of the transcriptional regulatory landscape of the developing mouse cortex identifies Neurog2 as a key epigenome remodeler. Nat Neurosci. 2022;25.

93. Zirkel A, Nikolic M, Sofiadis K, Mallm JP, Brackley CA, Gothe H, et al. HMGB2 Loss upon Senescence Entry Disrupts Genomic Organization and Induces CTCF Clustering across Cell Types. Mol Cell. 2018;70.

94. Divisato G, Chiariello AM, Esposito A, Zoppoli P, Zambelli F, Elia MA, et al. Hmga2 protein loss alters nuclear envelope and 3D chromatin structure. BMC Biol. 2022;20.

95. Wapinski OL, Vierbuchen T, Qu K, Lee QY, Chanda S, Fuentes DR, et al. Hierarchical mechanisms for direct reprogramming of fibroblasts to neurons. Cell [Internet]. 2013;155:621–35. Available from: http://www.ncbi.nlm.nih.gov/pubmed/24243019

96. Liu M-L, Zang T, Zou Y, Chang JC, Gibson JR, Huber KM, et al. Small molecules enable neurogenin 2 to efficiently convert human fibroblasts into cholinergic neurons. Nat Commun [Internet]. 2013;4. Available from: 10.1038/ncomms3183

97. Heinrich C, Bergami M, Gascón S, Lepier A, Dimou L, Sutor B, et al. Sox2-mediated conversion of NG2 glia into induced neurons in the injured adult cerebral cortex. Stem Cell Reports [Internet]. 2014;in press:1000–14. Available from: http://www.ncbi.nlm.nih.gov/pubmed/25458895

98. Liu ML, Zang T, Zhang CL. Direct Lineage Reprogramming Reveals Disease-Specific Phenotypes of Motor Neurons from Human ALS Patients. Cell Rep. 2016;14.

99. Kempf J, Knelles K, Hersbach BA, Petrik D, Riedemann T, Bednarova V, et al. Heterogeneity of neurons reprogrammed from spinal cord astrocytes by the proneural factors Ascl1 and Neurogenin2. Cell Rep. 2021;36.

100. Treutlein B, Lee QY, Camp JG, Mall M, Koh W, Shariati SA, et al. Dissecting direct reprogramming from fibroblast to neuron using single-cell RNA-seq. Nature [Internet]. 2016;534:391–5. Available from: https://www.ncbi.nlm.nih.gov/pubmed/27281220

101. Chen K, Zhang J, Liang F, Zhu Q, Cai S, Tong X, et al. HMGB2 orchestrates mitotic clonal expansion by binding to the promoter of C/EBPβ to facilitate adipogenesis. Cell Death Dis. 2021;12.

102. Ronfani L, Ferraguti M, Croci L, Ovitt CE, Schöler HR, Consalez GG, et al. Reduced fertility and spermatogenesis defects in mice lacking chromosomal protein Hmgb2. Development. 2001;128:1265–73.

103. Buffo A, Rite I, Tripathi P, Lepier A, Colak D, Horn AP, et al. Origin and progeny of reactive gliosis: A source of multipotent cells in the injured brain. Proc Natl Acad Sci U S A [Internet]. 2008/02/27. 2008;105:3581–6. Available from: http://www.ncbi.nlm.nih.gov/entrez/query.fcgi?cmd=Retrieve&db=PubMed&dopt=Citation&list_uids=18299565

104. Blum R, Heinrich C, Sanchez R, Lepier A, Gundelfinger ED, Berninger B, et al. Neuronal network formation from reprogrammed early postnatal rat cortical glial cells. Cereb Cortex [Internet]. 2010/06/22. 2011;21:413–24. Available from: http://www.ncbi.nlm.nih.gov/pubmed/20562320

105. Schindelin J, Arganda-Carreras I, Frise E, Kaynig V, Longair M, Pietzsch T, et al. Fiji: An open-source platform for biological-image analysis. Nat Methods. 2012.

106. Buenrostro JD, Wu B, Chang HY, Greenleaf WJ. ATAC-seq: A method for assaying chromatin accessibility genome-wide. Curr Protoc Mol Biol. 2015;2015.

107. Buenrostro JD, Giresi PG, Zaba LC, Chang HY, Greenleaf WJ. Transposition of native chromatin for fast and sensitive epigenomic profiling of open chromatin, DNA-binding proteins and nucleosome position. Nat Methods. 2013;10.

108. Quinlan AR, Hall IM. BEDTools: A flexible suite of utilities for comparing genomic features. Bioinformatics. 2010;26.

109. Ninkovic J, Pinto L, Petricca S, Lepier A, Sun J, Rieger MAA, et al. The transcription factor Pax6 regulates survival of dopaminergic olfactory bulb neurons via crystallin alphaA. Neuron [Internet]. 2010/11/26. 2010;68:682–94. Available from: http://www.ncbi.nlm.nih.gov/entrez/query.fcgi?cmd=Retrieve&db=PubMed&dopt=Citation&list_uids=21092858

110. Wiśniewski JR, Zougman A, Nagaraj N, Mann M. Universal sample preparation method for proteome analysis. Nat Methods. 2009;6:359–62.

111. Hauck SM, Dietter J, Kramer RL, Hofmaier F, Zipplies JK, Amann B, et al. Deciphering Membrane-Associated Molecular Processes in Target Tissue of Autoimmune Uveitis by Label-Free Quantitative Mass Spectrometry. Molecular & Cellular Proteomics. 2010;9:2292–305.

112. Merl J, Ueffing M, Hauck SM, von Toerne C. Direct comparison of MS-based label-free and SILAC quantitative proteome profiling strategies in primary retinal Müller cells. Proteomics. 2012;12:1902–11.

113. Ninkovic J, Steiner-Mezzadri A, Jawerka M, Akinci U, Masserdotti G, Petricca S, et al. The BAF Complex Interacts with Pax6 in Adult Neural Progenitors to Establish a Neurogenic Cross-Regulatory Transcriptional Network. Cell Stem Cell [Internet]. 2013;13. Available from: http://www.ncbi.nlm.nih.gov/pubmed/23933087

114. RStudio Team. RStudio: Integrated Development for R. RStudio, Inc., Boston, MA. URL http://www.rstudio.com/. RStudio, Inc. 2015;

