## Supplementary material for "Hmgb2 improves astrocyte to neuron conversion by increasing the chromatin accessibility of genes associated with neuronal maturation in a proneuronal factor-dependent manner": Suppl. Figures

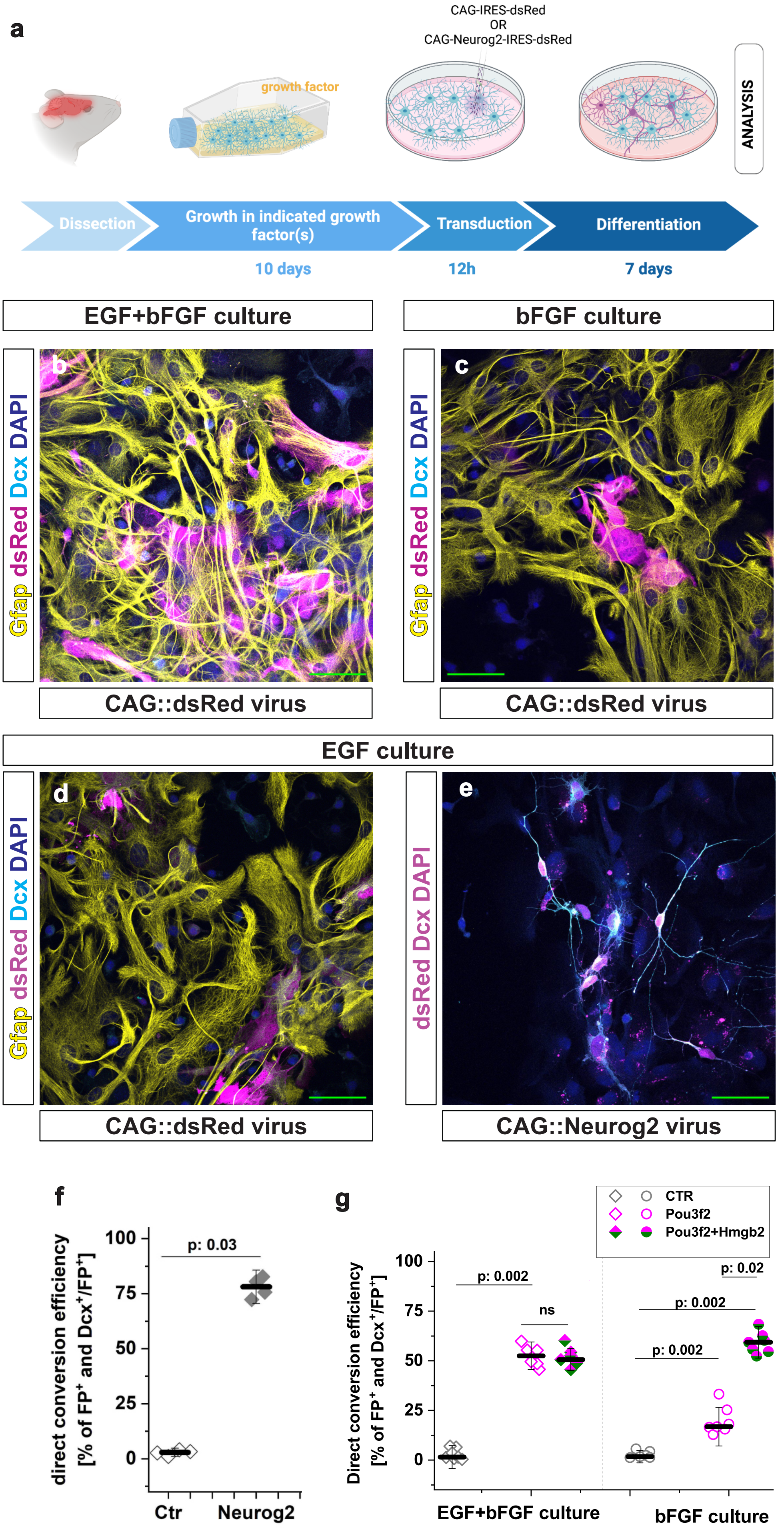


Maddhesiya, Lepko et al, Suppl. Figure 1

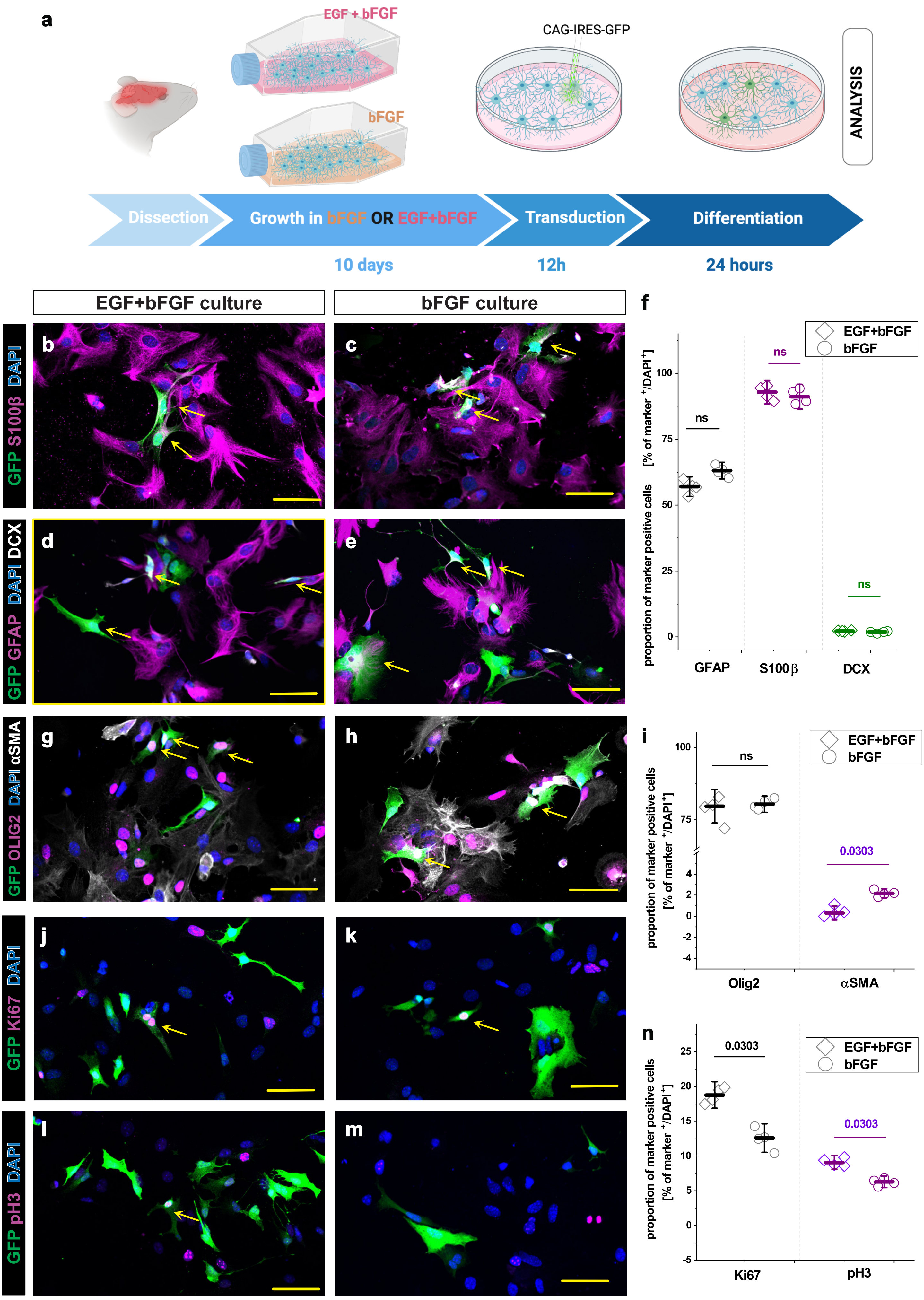


Maddhesiya, Lepko et al, Suppl. Figure 2

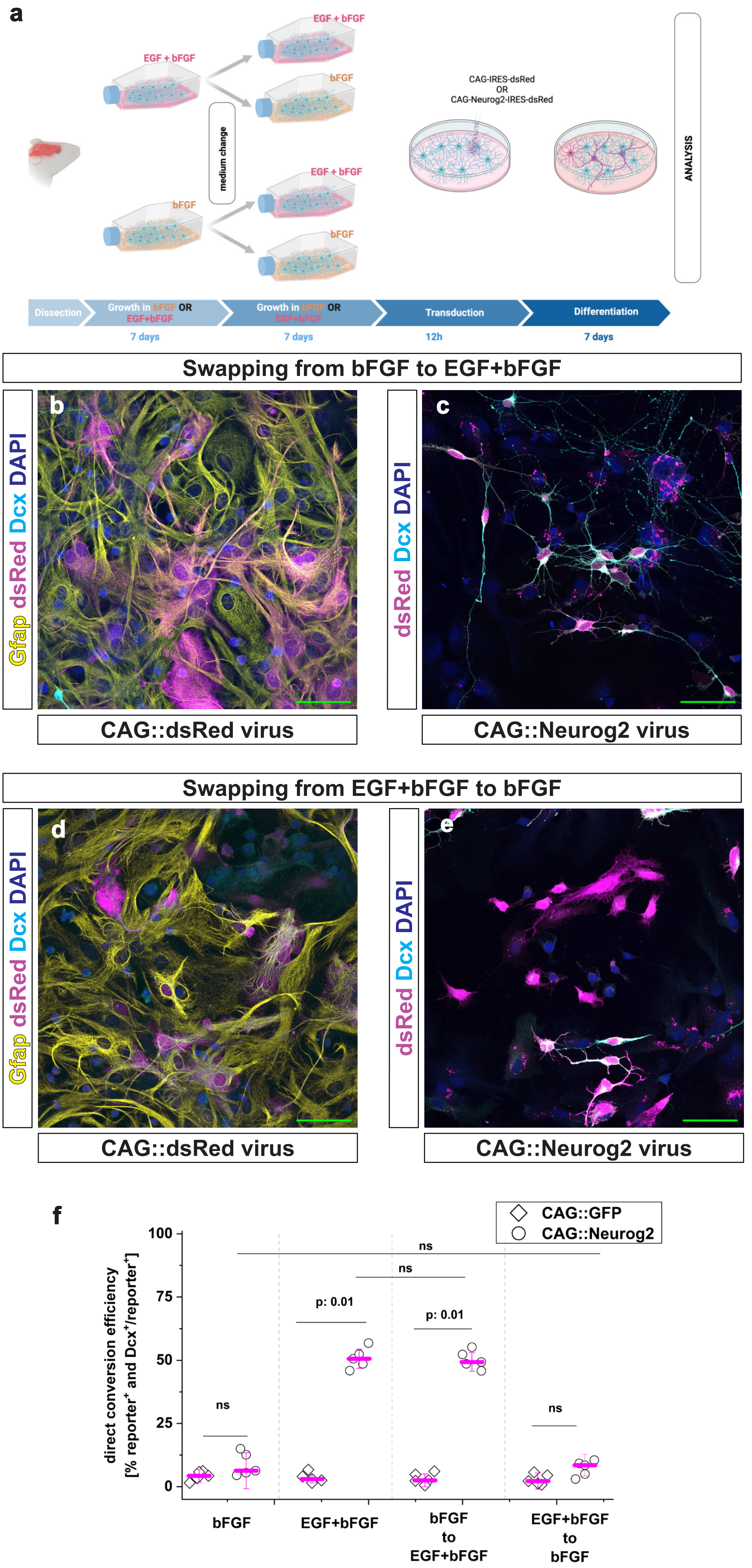


Maddhesiya, Lepko et al, Suppl. Figure 3

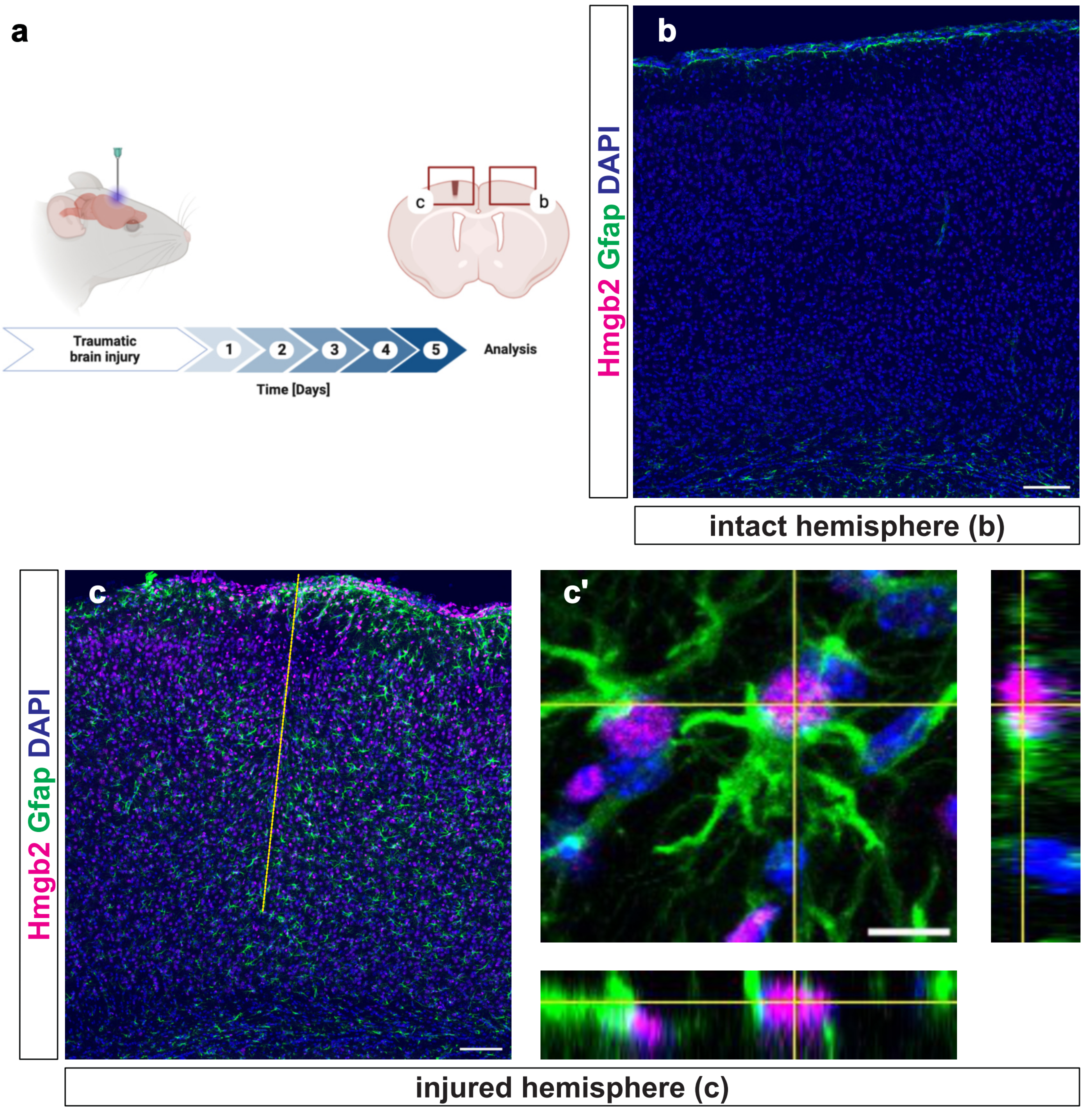


Maddhesiya, Lepko et al, Suppl. Figure 4

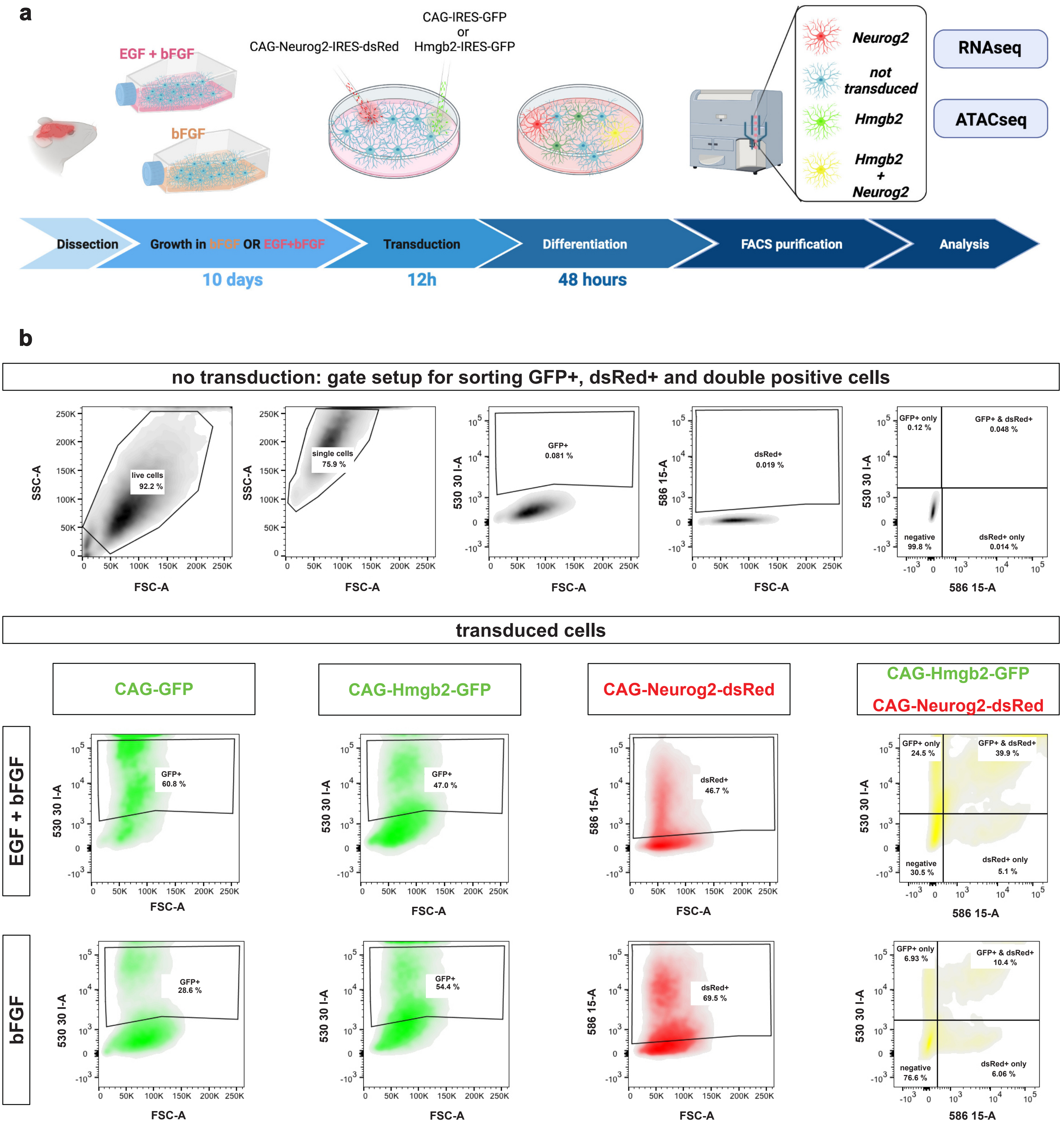


**Maddhesiya, Lepko et al, Suppl. Figure 5**

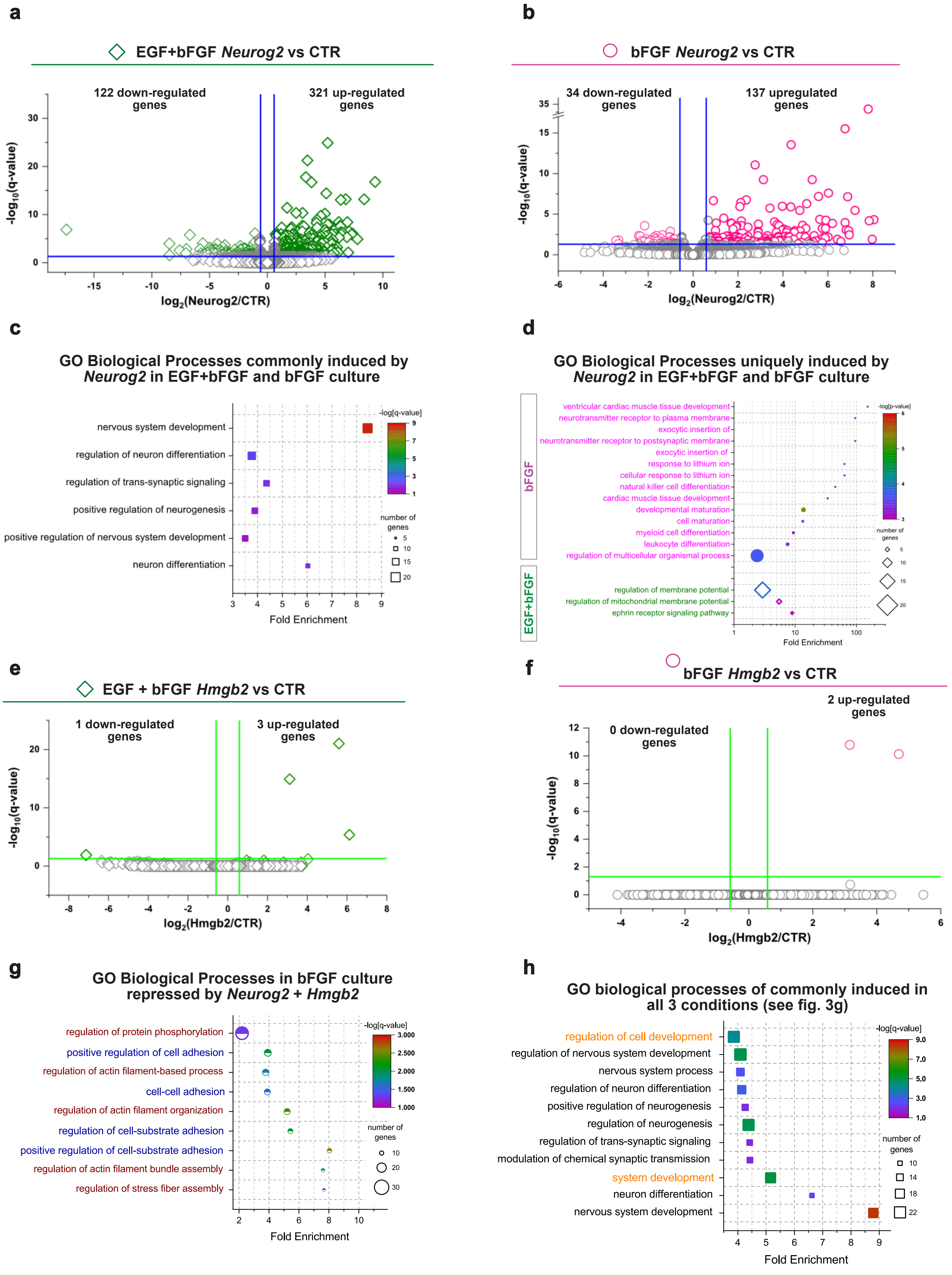


Maddhesiya, Lepko et al, Suppl. Figure 6

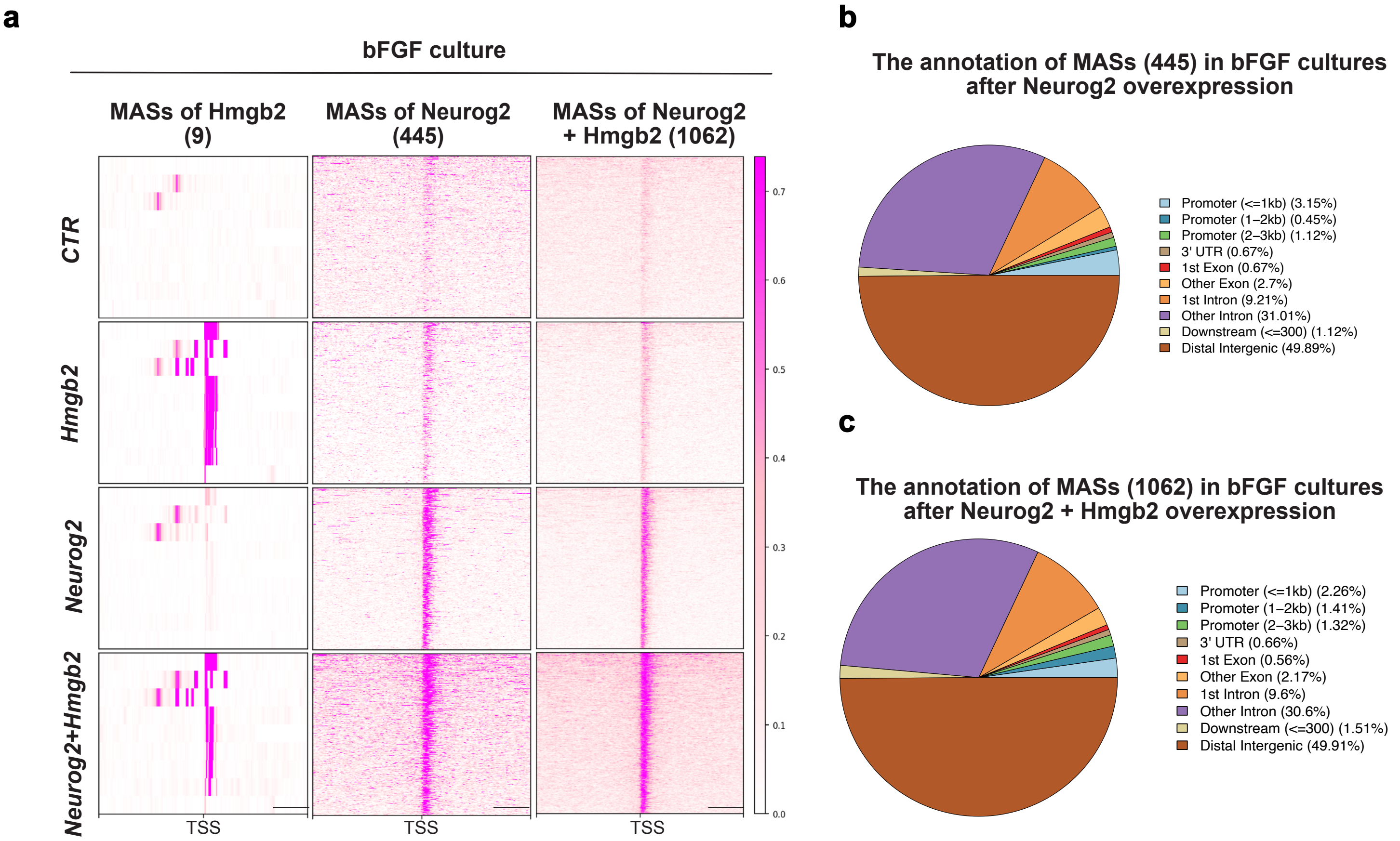


**Maddhesiya, Lepko et al, Suppl. Figure 7**

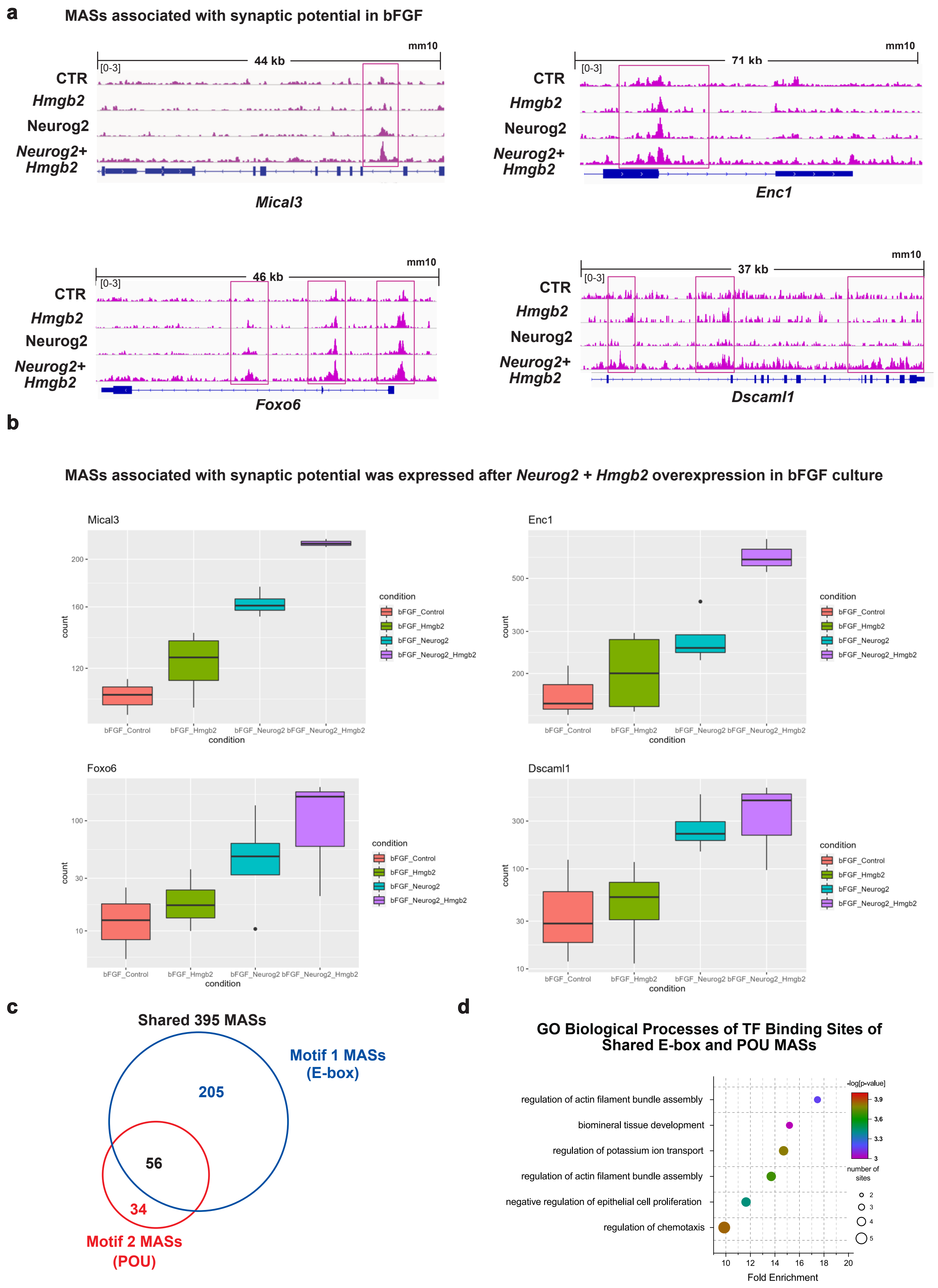


Maddhesiya, Lepko et al, Suppl. Figure 8
